# Certain Aquatic Eukaryotes Harbor an OLD-Like Immune Defense System

**DOI:** 10.64898/2026.08.18.745644

**Authors:** Halim Maaroufi

## Abstract

Key elements of eukaryotic antiviral immunity are evolutionarily conserved with prokaryotic anti-phage defense systems. The <u>O</u>vercoming <u>L</u>ysogenization <u>D</u>efect (OLD) anti-phage defense system is well characterized in prokaryotes but has remained unknown in eukaryotes. Here, the *old*-like genes are identified in certain aquatic eukaryotes, including the SAR supergroup, Filasterea, Chytridiomycota, and some invertebrate metazoan lineages (Placozoa, Cnidaria, Spiralia, Hemichordata, and Cephalochordata). Within molluscs, *old*-like genes are present in bivalves but absent in gastropods and cephalopods. Interestingly, the filasterean *Capsaspora owczarzaki*, an endosymbiont of the gastropod *Biomphalaria glabrata*, encodes putative secreted OLD-like proteins, suggesting it may provide its host with symbiont-mediated antiviral protection. Genomic and transcriptomic analyses revealed that *old*-like genes vary in copy and intron number and are expressed in some eukaryotic lineages. OLD-like proteins retain key structural features but exhibit a different topology from that of prokaryotes. Structural predictions reveal homodimeric dsDNA-bound architectures analogous to that of the *Bacillus cereus* VD045 GajA OLD homodimer. Furthermore, phylogenetic analysis divided OLD-like proteins into three distinct clusters, one of which likely represents an ancient inheritance from the ancestral Promethearchaeota (formerly ‘Asgard’ archaea) host. Together, these findings highlight an unexpected presence of *old*-like genes in certain aquatic eukaryotes, offering new insights into their evolutionary history and potential role in antiviral defense.

## 1. Introduction

For decades, researchers considered many immune mechanisms as evolutionary novelties of metazoans, but recent years several studies have indicated that eukaryotic key elements of cell-autonomous innate immune system trace back to prokaryotic anti-phage defense genes (Aravind et al., 2024; Litman et al., 2005). Multiple studies have demonstrated that key components of antiviral immunity are evolutionarily conserved between prokaryotes and eukaryotes, notably the Gasdermin, Viperin, and TIR-domain protein families (Johnson et al., 2022, Bernheim et al., 2021; Ofir et al., 2021). For example, in 2012 a major prokaryote signalling intermediate Cyclic di-(3′:5′)-guanosine monophosphate (c-di-GMP) was found in all major groups of Dictyostelia (Chen and Schaap, 2012). Many reviews detail the immune systems shared between prokaryotes and eukaryotes (Wein and Sorek, 2022; Yang et al., 2025).

The <u>O</u>vercoming <u>L</u>ysogenization <u>D</u>efect (OLD) anti-phage defense system is known in prokaryotes and some viruses (Akritidou and Thurtle-Schmidt, 2023). In 1970, the first OLD protein was identified in phage P2 as the element responsible for executing P2–Lambda interference (Lindahl et al., 1970). Class 1 OLD (P2 Old) system consists of single genes found in different genetic loci (Schiltz et al., 2019), whereas class 2 are composed of an OLD protein (GajA), and a UvrD/PcrA/Rep-like helicase (GajB) named Gabija defense system (Doron et al., 2018). Unlike the widespread Gabija system, Class 1 OLD family nucleases are present in only 0.45% of bacterial genomes but span a broad range of taxa (Patel and Seed, 2025). The OLD protein is composed of an N-terminal ABC-family ATPase domain associated with a C-terminal Toprim (topoisomerase-primase) nuclease domain. The structure of Class 1 OLD from *Thermus scotoductus* revealed another domain inserted into the ABC-family ATPase domain that allows the homodimerization of the OLD protein (Schiltz et al., 2020). Toprim is a divalent metal-binding domain found among other in <u>to</u>poisomerases and DnaG-type <u>prim</u>ases, hence the name Toprim. This domain is sufficient for nuclease activity and can cleave both circular and linear DNA. This cleavage follows a canonical divalent metal DNA cleavage mechanism.

The first structure of a full-length OLD protein was the Class 1 OLD in *T. scotoductus* that adopts a homodimeric assembly (Schiltz et al., 2020). The structure of class 2 OLD (GajA) adopts a homotetramer architecture. GajB binds on the GajA tetramer to form a multimeric GajA-GajB assembly (Antine et al., 2024). In this study, *old*-like genes were identified in certain aquatic eukaryotes. Phylogenetically, they constitute three distinct clusters, one of which likely represents an ancient inheritance from the ancestral Promethearchaeota (formerly ‘Asgard’ archaea) host. Structural predictions further reveal that eukaryotic OLD-like homologues adopt an overall architecture closely resembling the prokaryotic Class 1 and 2 (GajA) OLD nucleases. These findings show an unexpected diversity of eukaryotic OLD-like proteins, shedding light on their evolutionary history and providing a potential new gene resource for antiviral defense systems.

## 2. Materials and methods

### 2.1. Identification of probable eukaryotic OLD-like proteins

To identify Overcoming Lysogenization Defect (OLD)-like protein sequences across eukaryotes, homology searches were executed using the NCBI BLAST web server. Initial queries comprised the well-characterized prokaryotic OLD-family defense proteins *Bacillus cereus* VD045 GajA OLD (UniProt ID: J8H9C1) and *Thermus scotoductus* OLD (UniProt ID: E8PLM2). Protein-level searches were conducted by DELTA-BLAST against the eukaryotic Clustered NR protein database (nr_clustered_seq). Complementary nucleotide-level mapping was performed by TBLASTN against both the Core Nucleotide database (core_nt) and the Whole Genome Shotgun database (WGS).

To filter out potential prokaryotic contaminants and validate true eukaryotic candidates, a reverse BLAST strategy was performed. All retrieved eukaryotic OLD-like candidate sequences were subjected to reverse BLASTP searches against the prokaryotic subsets of the nr_clustered_seq database, followed by reverse TBLASTN searches against the prokaryotic partitions of the core_nt and WGS databases. Candidate sequences displaying higher similarity to prokaryotes than to eukaryotic lineages were classified as contaminants and excluded from downstream analyses.

### 2.2. Intron analysis of eukaryotic old-like genes

To determine whether the identified eukaryotic sequences contain introns, their genomic architecture was resolved by mapping the TBLASTN alignments against the raw genomic chromosomes and scaffolds within the WGS database.

### 2.3. Structural search of probable OLD-like proteins in Eukaryotes

To discover potential OLD-like proteins across diverse eukaryotic taxonomic groups, structure-based searches were conducted using Foldseek (van Kempen et al., 2024) and DALI (Holm et al., 2023) programs. The structures of *B. cereus* VD045 GajA OLD (PDB ID: 8JQ9) and *T. scotoductus* OLD (PDB ID: 6P74) were used as queries against the Protein Data Bank (PDB) (Vallat et al., 2026) and AlphaFold (Varadi et al., 2024) databases.

### 2.4. Domain prediction of OLD-like proteins

Domain architecture annotation of the retrieved protein sequences was performed using hmmscan on the HMMER web server (Rajkovic et al., 2026) and InterProscan (https://www.ebi.ac.uk/interpro/). HMMER web server, protein sequences were queried against the <u>Pfam database</u> (v37.2) using the default gathering threshold as the score cutoff. Furthermore, to define protein domain architecture, the NCBI CDD-Search (https://www.ncbi.nlm.nih.gov/Structure/cdd/wrpsb.cgi) was used (Wang et al., 2023). The parameters of search are: Pfam-19638 PSSMs database, cutoff is 0.01 and the result mode is concise.

### 2.5. Evolutionary analysis of OLD-like proteins

To elucidate the evolutionary history of the OLD-like proteins across prokaryotic and eukaryotic lineages, phylogenetic reconstruction was performed using a maximum-likelihood framework.

Multiple sequence alignment of the 32 protein sequences was carried out using MAFFT v7, utilizing the high-accuracy iterative refinement method L-INS-i (--localpair --maxiterate 1000) to optimize local homologies among highly divergent taxa (Katoh and Standley, 2013). To minimize phylogenetic noise and eliminate uninformative alignment regions, automated trimming was performed using TrimAl v1.4 with the heuristically optimized-automated algorithm (Capella-Gutiérrez et al., 2009). This procedure yielded a clean, highly informative alignment blueprint of 213 columns, of which 205 were parsimony-informative. Prior to tree inference, the alignment was subjected to a chi-squared (*χ*2) test of sequence composition implemented in IQ-TREE v2.1.3 to ensure composition stationarity; 25 out of 32 taxa successfully passed the test (*p*>0.05, *df*=19) (Minh et al., 2020).

Phylogenetic tree inference was performed using IQ-TREE v2.1.3. To effectively mitigate potential systematic errors, such as long-branch attraction (LBA) and compositional heterogeneity typical of deep-scale or horizontal gene transfer (HGT) datasets, the advanced site-heterogeneous mixture model LG+C20+F+G was employed (Le et al., 2008). Branch supports were thoroughly evaluated using 1000 replicates of the UltraFast Bootstrap approximation (UFBoot) (Hoang et al., 2018). The tree was inferred without a pre-computed guide tree to ensure unbiased site-frequency estimation by the mixture model. Given the uncharacterized evolutionary origin of the eukaryotic OLD-like sequences and the potential presence of horizontal gene transfers, the final tree topology was rooted using the midpoint rooting method to avoid *a priori* evolutionary assumptions. The final phylogenetic tree was midpoint-rooted and visualized using FigTree (v1.4.3) (http://tree.bio.ed.ac.uk/software/figtree/). In addition, multiple sequence alignment obtained by MAFFT v7 was visualized with ENDscript (Gouet et al., 2002).

### 2.6. Structure prediction of OLD-like proteins

AlphaFold 3 (Abramson et al., 2024) was used to predict the structures of apo OLD-like proteins (monomer and homodimer) and OLD-like proteins (homodimer) complexed to different combinations of ions (Mg^2+^ and Ca^2+^), ATP and a 21-bp dsDNA substrate (PDB ID: 8X51). Electrostatic potential surface mapping of OLD-like proteins was conducted on the webserver: https://server.poissonboltzmann.org/pdb2pqr.

Topology diagrams of 3D structure of monomers are generated by PDBsum1 (Laskowski, 2022). All structural figures were prepared with PyMOL v2.3.2.

### 2.7. Prediction of secretory OLD-like proteins from Capsaspora

To determine whether any OLD-like proteins from *Capsaspora* are target proteins for classical secretion, sequence-based analyses were performed using SignalP v3.0 (Bendtsen et al., 2004) and Phobius (Käll et al., 2004). Phobius was specifically employed to effectively differentiate between hydrophobic N-terminal signal peptides and downstream transmembrane helices.

## 3. Results and discussion

### 3.1. OLD-like proteins in eukaryotes

The <u>O</u>vercoming <u>L</u>ysogenization <u>D</u>efect (OLD) anti-phage defense system is well-characterized in prokaryotes and certain viruses (Dot et al., 2023). In this study, *old*-like genes were identified in Sar supergroup (protists), Chytridiomycetes (Primitive flagellated fungi), Filasterea (Unicellular amoeboid organisms), Placozoa (’flat animals’), Cnidaria (including Octocorallia [soft corals], Scleractinia [stony corals], and Actiniaria [sea anemones]), Spiralia (bivalve molluscs, Rotifera, and Nemertea), Hemichordata, and Cephalochordata (*Branchiostoma* and *Asymmetron*) (Table S1).

These taxonomic groups represent a broad evolutionary spectrum. The Sar supergroup and Filasterea consist of unicellular organisms. Among them, Filasterea are close evolutionary relatives of Metazoa that possess genetic precursors for multicellularity, such as cell-to-cell adhesion genes (Jacques et al., 2022; Li et al., 2025). Predominantly unicellular, Chytridiomycetes (known as chytrids) are aquatic, primitive, flagellated fungi.

In multicellular invertebrate lineages, Placozoa is a primitive phylum comprising the simplest free-living marine invertebrates (Schierwater and DeSalle, 2018). However, the placozoan *Trichoplax* has a diversity of immune-related genes (Romanova and Moroz, 2025). Whereas Corals and sea anemones are diploblastic animals exhibiting radial symmetry and a rudimentary nervous system. The more derived Rotifera, Nemertea and bivalve molluscs are triploblastic and bilaterally symmetrical organisms that possess specialized organs and a centralized nervous system (Dunn et al., 2008). Within the deuterostomes, Hemichordata share a close evolutionary link to vertebrates (Chordates) and have dorsal and ventral nerve cords. Finally, Cephalochordata represents a subphylum of small, fish-like marine chordates, where *Branchiostoma lanceolatum* serves as a model organism for understanding the evolutionary transition from early chordates to complex vertebrates.

A reverse BLAST strategy confirmed that these eukaryotic OLD-like sequences are authentic genomic elements rather than prokaryotic contaminants (see § 2.1). Furthermore, eukaryotic OLD-like proteins share low sequence identity with their prokaryotic homologues. For example, the OLD nuclease-like protein of the bivalve mollusc *Magallana gigas* (UniProt ID: A0A8W8L770) exhibits only 13% amino acid identity (61/457) and 30% similarity (138/457) to the Class 1 OLD nuclease of *T. scotoductus* (UniProt ID: E8PLM2). Similarly, it displays 14% identity (63/462) and 28% similarity (130/462) to GajA from *B. cereus* VD045 (UniProt ID: J8H9C1). In addition, scanning the OLD-like sequences by the NCBI CDD-Search, HMMER and InterProscan web servers confirm that they contain the ABC-family ATPase and TOPRIM domains. The amino acid sequence length of OLD-like proteins ranges from 577 to 844 amino acids (Table S2), and DeepLoc 2.1 (Ødum et al., 2024) predicts that they are cytoplasmic.

Multiple sequence alignments of these aquatic eukaryotic OLD-like proteins revealed that the core amino acids motifs (Walker A motif (for ATP binding), Walker B motif (for ATP hydrolysis), and catalytic residues in the nuclease domain) found in the Class 1 OLD nuclease of *T. scotoductus* and GajA of *B. cereus* VD045 remain conserved, suggesting that OLD-like proteins are probably enzymatically active (**Fig. 1**). However, lineage-specific variations occur within the functional D-loop. The catalytic histidine residue that replaces the conserved aspartic acid in class 1 OLD, is universally conserved in most eukaryotic OLD-like proteins, but it is substituted in *Capsaspora owczarzaki* (Filasterea), *Adineta* (Rotifera), and *Cephalothrix simula* (ribbon worms) by a tyrosine, phenylalanine and asparagine, respectively (**Fig. S1**).

**Fig. 1.**
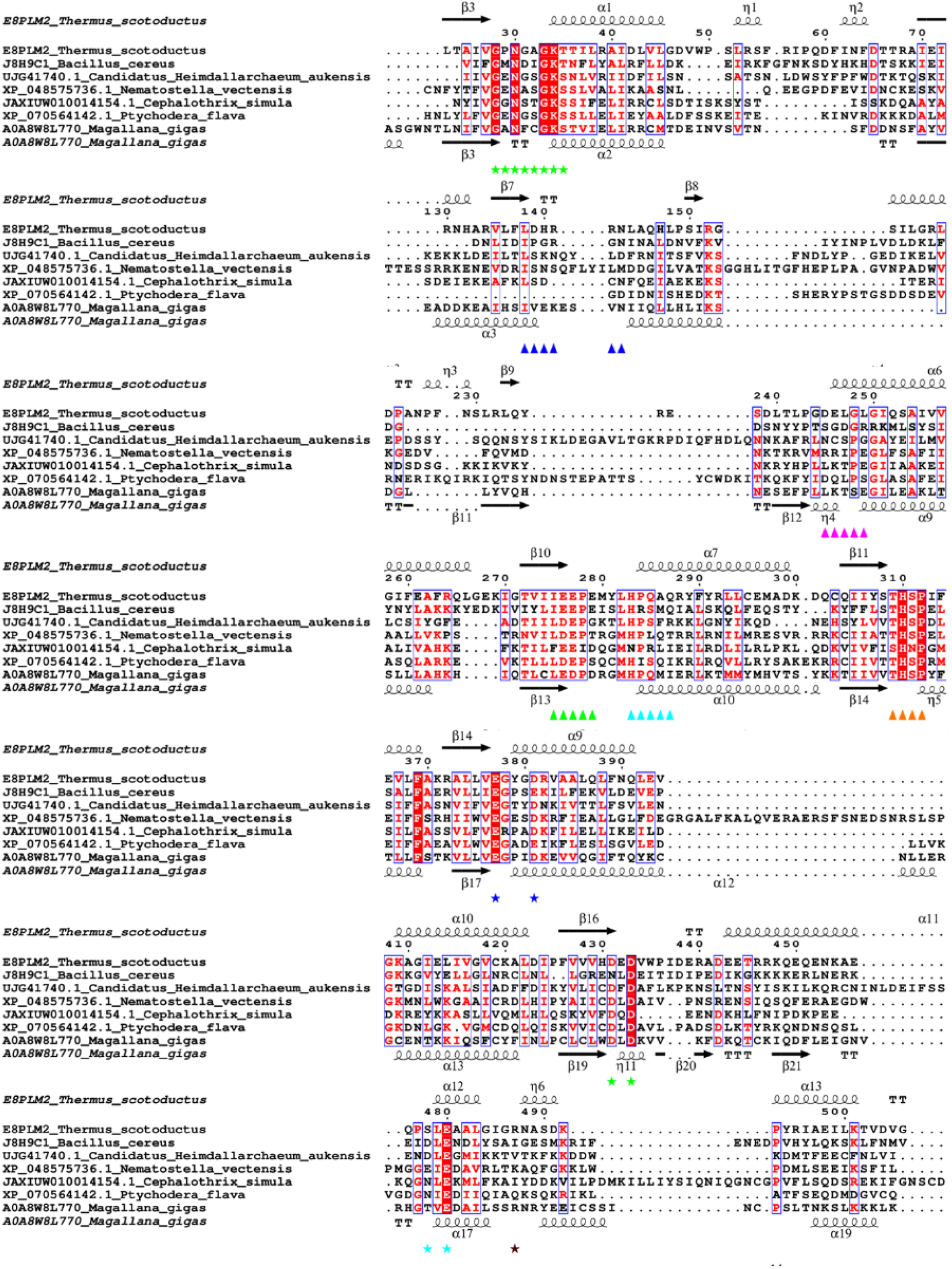
Multiple sequence alignment of some OLD-like proteins from aquatic eukaryotes. Conserved functional motifs of *T. scotoductus* class 1 OLD are indicated by distinct symbols and colors: Walker A (P-loop) (green stars), Q-loop (blue triangles), signature sequence (pink triangles), Walker B (green triangles), D-loop (cyan triangles), H-loop (orange triangles), Metal A and Metal B motifs (blue stars), Metal A motif (D-x-D, green stars), Metal B motif (S-x-E, cyan stars) and spatially conserved basic residue R487 residue (brown star) required for OLD nuclease cleavage. Secondary structures of *T. scotoductus* (PDB ID: 6P74) and *M. gigas* (predicted by AF3) are indicated above and below MSA, respectively. Taxonomy of species in MSA, *T. scotoductus* and *B. cereus* VD045 (Bacteria), *Candidatus Heimdallarchaeum aukensis* (Archaea), *Nematostella vectensis* (Cnidaria), *Cephalothrix simula* (ribbon worms), *Ptychodera flava* (Hemichordata), and *M. gigas* (Bivalvia). Accession numbers for all sequences are listed preceding the respective species names. Only sequences regions containing the relevant amino acid motifs are displayed.

In prokaryotes, the Gabija defense system operates as a heteromeric complex consisting of both GajA and GajB proteins (Doron et al., 2018). While GajA contains core ABC-family ATPase and Toprim domains structurally similar to the OLD protein. In this study, sequence and structural searches for eukaryotic GajB homologues yielded no matches.

### 3.2. Phylogenetic analysis

Phylogenetic analysis showed that eukaryotic OLD-like proteins split into three distinct clusters, with the first two clusters displaying robust bootstrap support (**Fig. 2**). Cluster 1, comprising Promethearchaeota (Imachi et al., 2024; Vosseberg et al., 2024) and Metazoa, represents a lineage derived from the ancestral archaeal host cell involved in eukaryogenesis (Tobiasson et al., 2026) and the contribution to the origins of antiviral defense systems in eukaryotes (Leão et al., 2024). However, its restriction to specific metazoan groups, such as Cnidaria and select bilaterians like bivalve molluscs and ribbon worms, suggests a complex evolutionary history. This pattern could be explained by an ancient vertical inheritance marked by subsequent extensive gene loss, or alternatively, by an early horizontal gene transfer (HGT) acquisition event captured by the common ancestor of these specific animal phyla. This specific metazoan grouping aligns well with recent chromosome-scale synteny analyses, which demonstrate that cnidarians and bilaterians share rare, irreversible ancestral chromosome fusion-and-mixing events, which could have trapped or stabilized a horizontally acquired or vertically inherited defense module (Callaway, 2026; Schultz et al., 2023). Conversely, Cluster 2, comprising SAR supergroup, *Candidatus* Sungbacteria (Patescibacteria), and *Candidatus* Hodarchaeales (Promethearchaeota), demonstrates an independent HGT event distinct from Cluster 1. Lastly, Cluster 3 forms an intermixed assemblage of prokaryotic and eukaryotic lineages. Due to low internal bootstrap support, the precise evolutionary trajectories and HGT directions within this cluster remain unresolved.

**Fig. 2.**
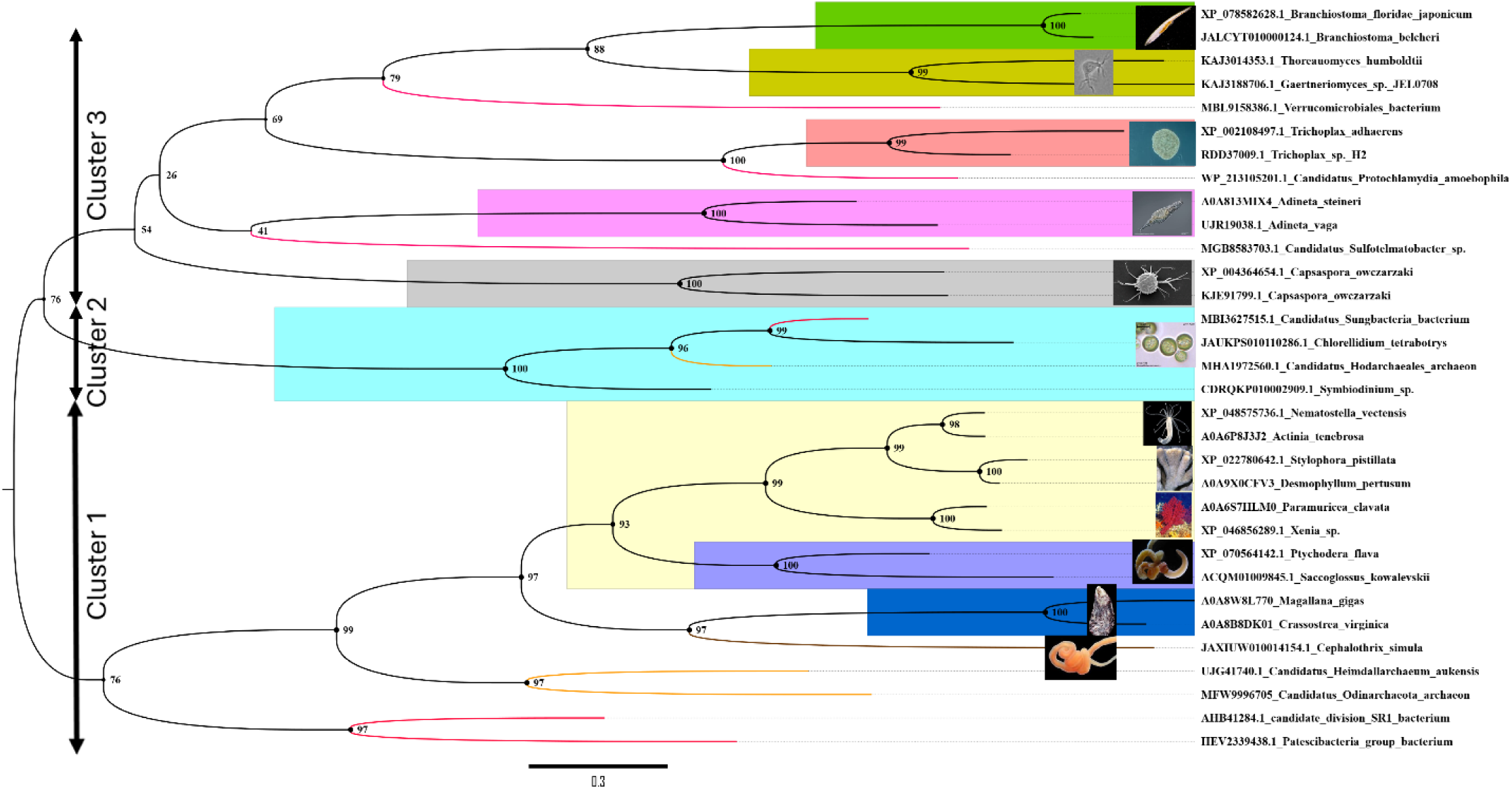
Midpoint-rooted phylogenetic tree of eukaryotic OLD-like proteins. Lineages are color-coded as follows: Bacteria (red branches), Archaea (orange branches), SAR supergroup (cyan), Filasterea (grey), Chytridiomycota (olive green), and metazoan lineages: Placozoa (Light Salmon Pink), Cnidaria (yellow), Spiralia clade is represented by Nemertea (brown branch), Bivalvia (blue), and Rotifera (pink), Hemichordata (purple), and Cephalochordata (green). The maximum-likelihood tree topology was inferred using IQ-TREE and visualized using FigTree v1.4.3.

Our phylogenetic analysis uncovers a split evolutionary origin for the *old*-like gene that mirrors the deepest divides in the eukaryotic tree of life. Within Chytridiomycetes, Filasterea, and all examined Metazoa, the *old*-like gene clusters robustly with homologs from Promethearchaeota, indicating likely vertical inheritance dating back to eukaryogenesis. Conversely, in the distantly related SAR supergroup, the gene appears to be the product of an independent horizontal gene transfer (HGT) event from a prokaryotic donor.

### 3.3. Capsaspora and gastropod molluscs

While OLD-like proteins were highly prevalent in bivalve molluscs, they were entirely absent in the sequences and genomic datasets of gastropods and cephalopods (**Table S1**). Of note, during a structural and sequence analysis to classify the ABC ATPase superfamily, Krishnan et al. reported that the OLD family underwent independent transfers to the unicellular holozoan *C. owczarzaki* and the bivalve mollusc *Crassostrea* (Krishnan et al., 2020). The Filasterean *C. owczarzaki*, one of the closest unicellular relatives of multicellular animals, is an endosymbiont found in the hemolymph of the gastropod *Biomphalaria glabrata* (Hertel et al., 2002). *C. owczarzaki* genome encodes 13 OLD-like proteins. Interestingly, two of which (GenBank IDs: KJE94168.1 and KJE88441.1) were predicted by Phobius and SignalP (v3.0) to possess an N-terminal signal peptide (Sperschneider et al., 2015). This finding suggests that these two proteins are likely secreted in the hemolymph of the gastropod. Consequently, the host gastropod may passively benefit from this putative antiviral defense system; reflecting a mutualistic cooperation between these symbiotic partners (Galvão Ferrarini et al., 2025).

### 3.4. Genomic and transcriptomic analyses of old-like genes

Comparative genomic architecture in eukaryotes containing the *old*-like gene revealed highly variable, taxon-dependent profiles of intron distribution (**Table S3**). Single-celled protists (*Symbiodinium sp*., and *Chlorellidium tetrabotrys*) within the SAR supergroup exclusively contained a single, intronless *old*-like gene, indicating a high probability of HGT from prokaryotic sources in this lineage. Unicellular Placozoa and Chytridiomycetes also contain intronless *old*-like gene. In contrast, diverse animals, including Cnidaria (sea anemones, stony corals, and soft corals) and Spiralia (Nemertea and Rotifera) possesses only a single gene with or without intron. Interestingly, in the unicellular holozoan *Capsaspora owczarzaki* (Filasterea) genome functions as a dynamic genetic mosaic, retaining entirely uninterrupted coding regions on chromosomes 2 and 5, while simultaneously housing heavily modified variants containing up to 5 introns on chromosome 1. A similarly complex intragenomic architecture is mirrors within the bivalve mollusc *Magallana gigas*, which demonstrates extensive copy number expansion and structural heterogeneity across multiple chromosomes. Within the *M. gigas* genome, chromosome 3 harbors a cluster comprising three intronless loci alongside one intron-containing gene variant, while chromosome 6 houses an even split of two intronless genes and two variants containing two introns. Additional copies are distributed on chromosome 7, which contains two intronless variants and two genes with probably introns, as well as a solitary intronless copy on chromosome 9. When compared to the architectural layout observed in *C. owczarzaki*, *M. gigas* exhibits a fundamentally different pattern of genomic expansion. While *C. owczarzaki* clusters its structural variants distinctly by chromosome, segregating fully continuous genes on separate chromosomes away from highly intronized loci, *M. gigas* presents a highly duplicated, mixed mosaic configuration. Within individual chromosomal loci (such as chromosomes 3 and 6), *M. gigas* co-localizes ancestral-state intronless genes immediately alongside multi-exon configurations, signaling localized duplication events and highly active, variable intronization mechanisms acting unevenly within the same chromosomal neighborhoods. Taken together, these results demonstrate that the structural organization of *old*-like genes varies non-uniformly in eukaryotic genomes, transitioning from probable recent HGT-driven intronless configurations in SAR supergroup to highly complex, variable multi-exon architectures in metazoans (Wei et al., 2023). Furthermore, transcripts of *old*-like genes from Placozoa, sea anemones, stony and soft corals, bivalves, Hemichordata, and Cephalochordata are found in transcriptomes of NCBI database (**Table S3**).

### 3.5. Quaternary structure prediction of M. gigas OLD-like protein

AlphaFold 3 (AF3) was used to predict the structures of the *M. gigas* OLD-like protein (MgOLD-like) under various conditions. These include the apo monomer, the apo homodimer, homodimers bound to individual ligands (Mg²⁺, Ca²⁺, ATP, or double-stranded DNA (dsDNA)), and homodimers bound to various combinations of these ligands (**Table S4**).

The monomer structure of *B. cereus* GajA (BcGajA) (PDB ID: 8SM3) consists of 21 β-strands and 21 α-helices, while the OLD *T. scotoductus* (TsOLD) (PDB ID: 6P74) is composed of 17 β-strands and 14 α-helices (Antine et al., 2023; Schiltz et al., 2020). In comparison, the MgOLD-like contains 21 β-strands and 21 α-helices (**Fig. S2**). Its ATPase domain core consists of 15 β-strands and 10 α-helices, whereas its Toprim domain contains 6-stranded parallel β-sheet surrounded by 8 α-helices (**Fig. 3** and **Fig. S2**). The dimerization domain is formed with amino acids between 213 and 277, and the extra helical domain insert encompasses residues 555-606. This latter domain is present in Class 2 OLD but absent in Class 1 OLD (Schiltz et al., 2020; Antine et al., 2023; Li et al., 2024). Notably, the conformation of the extra helical domain insert shifts upon dsDNA binding. Compared to the architecture of class 1 TsOLD and class 2 BcGajA, MgOLD-like monomer retains key structural features (ATPase /TOPRIM domains and catalytic active sites) but exhibit a distinct topology from that of prokaryotes (**Fig. S2**). Indeed, structural superposition of MgOLD-like monomer with TsOLD and BcGajA monomers yields an RMSD of 4.977 Å (over 152 amino acids) and 5.616 Å (over 192 amino acids), respectively.

**Fig. 3.**
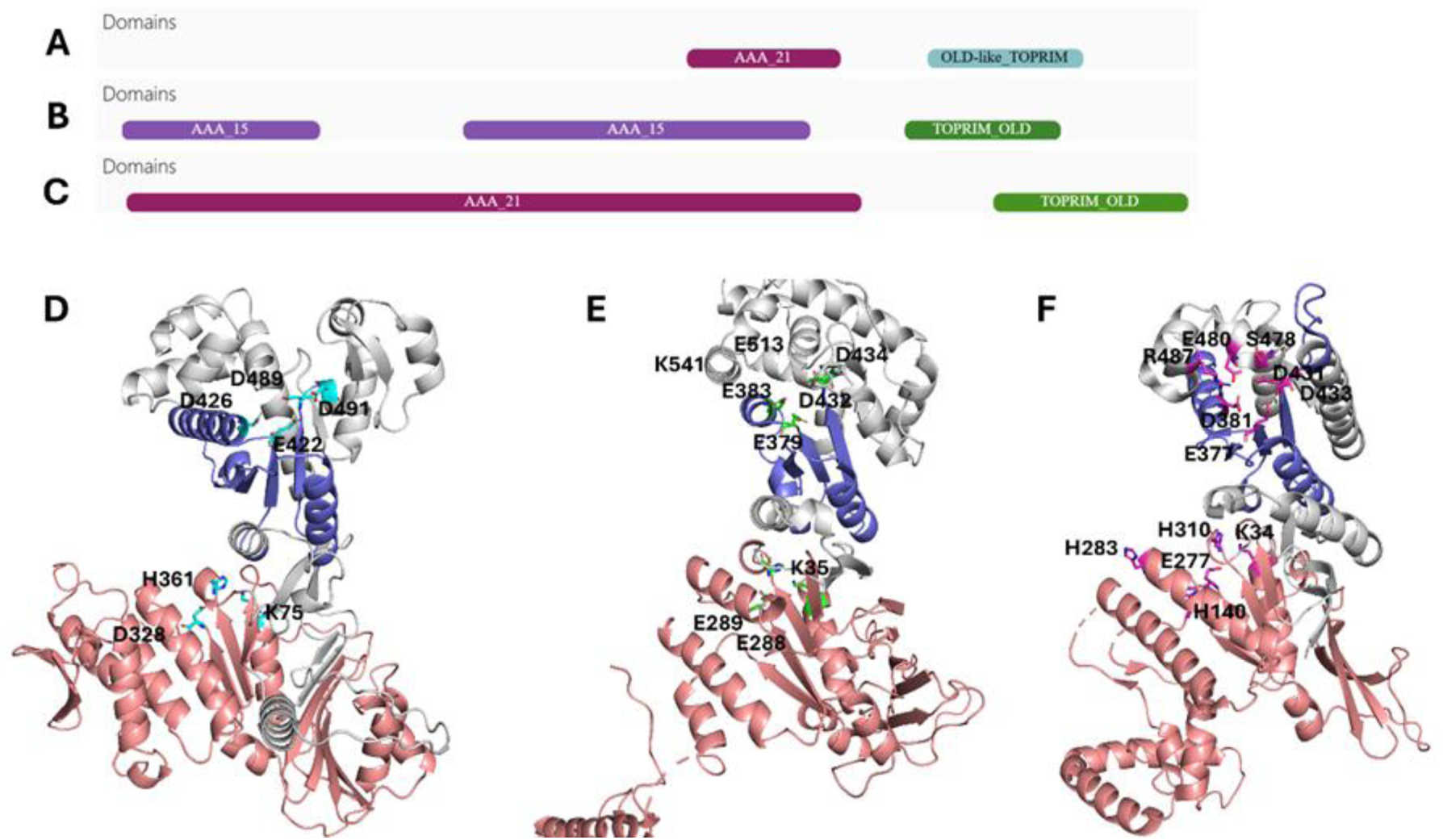
Comparative structural architecture of *M. gigas* OLD-like, Class 2 *B. cereus* GajA, and Class 1 *T. scotoductus* monomers. The N-terminal ATPase domain is colored salmon, and the C-terminal Toprim domain is colored purple. (**A–C**) Schematic domain architectures of (**A**) *M. gigas* OLD-like protein, (**B**) *B. cereus VD045* GajA, and (**C**) *T. scotoductus* OLD protein. (**D– F**) Predicted monomer structures of (**D**) *M. gigas* OLD-like protein, (**E**) Class 2 OLD monomer from *B. cereus VD045* GajA (PDB ID: 8SM3), and (**F**) Class 1 OLD monomer from *T. scotoductus* (PDB ID: 6P74). Active site residues within both the ATPase and Toprim domains are highlighted in stick representation.

OLD-family nucleases typically use a two-metal-ion catalytic mechanism for DNA cleavage (Akritidou and Thurtle-Schmidt, 2023; Dot et al., 2023). AF3 modeling predicted that apo MgOLD-like proteins preferentially assemble into homodimers with high confidence scores (pTM: 0.79, ipTM: 0.77) (**Table S4**). Conversely, alternative homotetrameric configurations yielded significantly lower scores (pTM: 0.50, ipTM: 0.40), suggesting that the functional unit of MgOLD-like is a homodimer. Structural alignment of the apo MgOLD-like homodimer with TsOLD and BcGajA homodimers revealed substantial structural divergence, resulting in large RMSDs of 5.75 Å (over only 248 amino acids) and 6.35 Å (over only 208 amino acids), respectively.

In ligand-bound AF3 simulations, Ca^2+^ ions and ATP consistently docked into their predicted canonical pockets within the MgOLD-like homodimer (**Fig. 4** and **Fig. 5**). Interestingly, Mg^2+^ ions showed a propensity to mislocalize to the Ca^2+^ binding site unless co-modeled with ATP, or within the fully assembled quaternary complex containing Ca^2+^, ATP, and dsDNA (**Table S4**). In this complete, fully coordinated complex, the functional homodimer architecture coordinates different ligands across distinct domains (**Fig. 4** and **Fig. 5A**). Mg^2+^ ions are localized at the ATPase active site of each monomer and coordinated by the phosphates of the ATP and three residues, K76, D328, and H361. Whereas the 21-bp dsDNA fragment is tightly sandwiched at the top by the symmetric Toprim domains. The Toprim nuclease active site, a cluster of conserved acidic residues comprising E422, D426, D489, and D491 forms the catalytic core, where they directly coordinate Ca^2+^ ions essential for DNA nuclease activity.

**Fig. 4.**
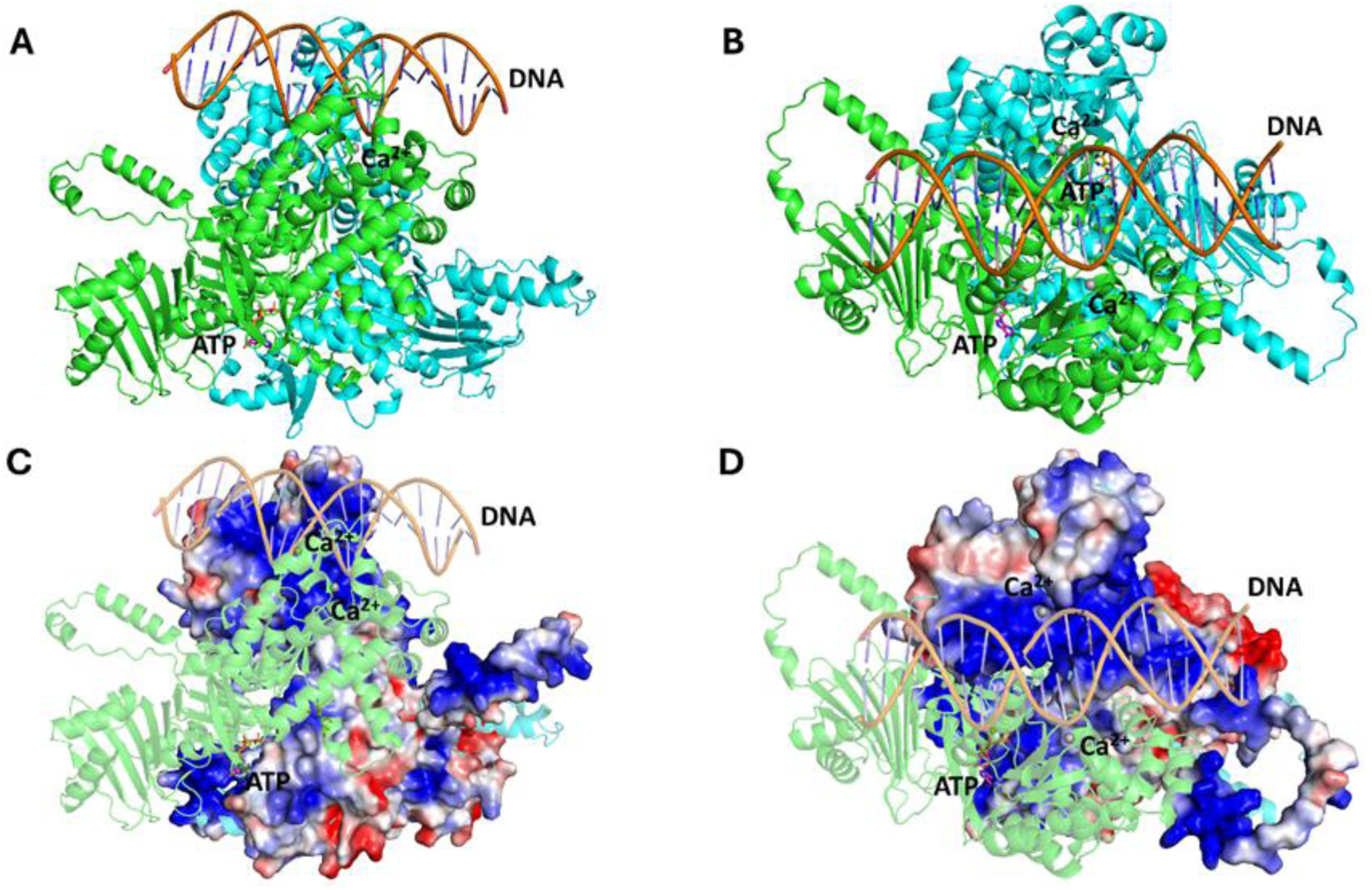
Predicted homodimer models of the *M. gigas* OLD-like protein bound to Ca²⁺, Mg²⁺, ATP, and dsDNA. (**A**) Front and (**B**) top views of the *M. gigas* homodimer. Monomer A is in green, monomer B is cyan, dsDNA is orange, ATP is light gray, Ca²⁺ ions are gray spheres, and Mg²⁺ ions are yellow-orange spheres. (**C**) Side and (**D**) top views of monomer B showing the electrostatic potential surface mapped onto the cartoon representation of the homodimer. Negative and positive electrostatic potentials are colored red and blue, respectively.

**Fig. 5.**
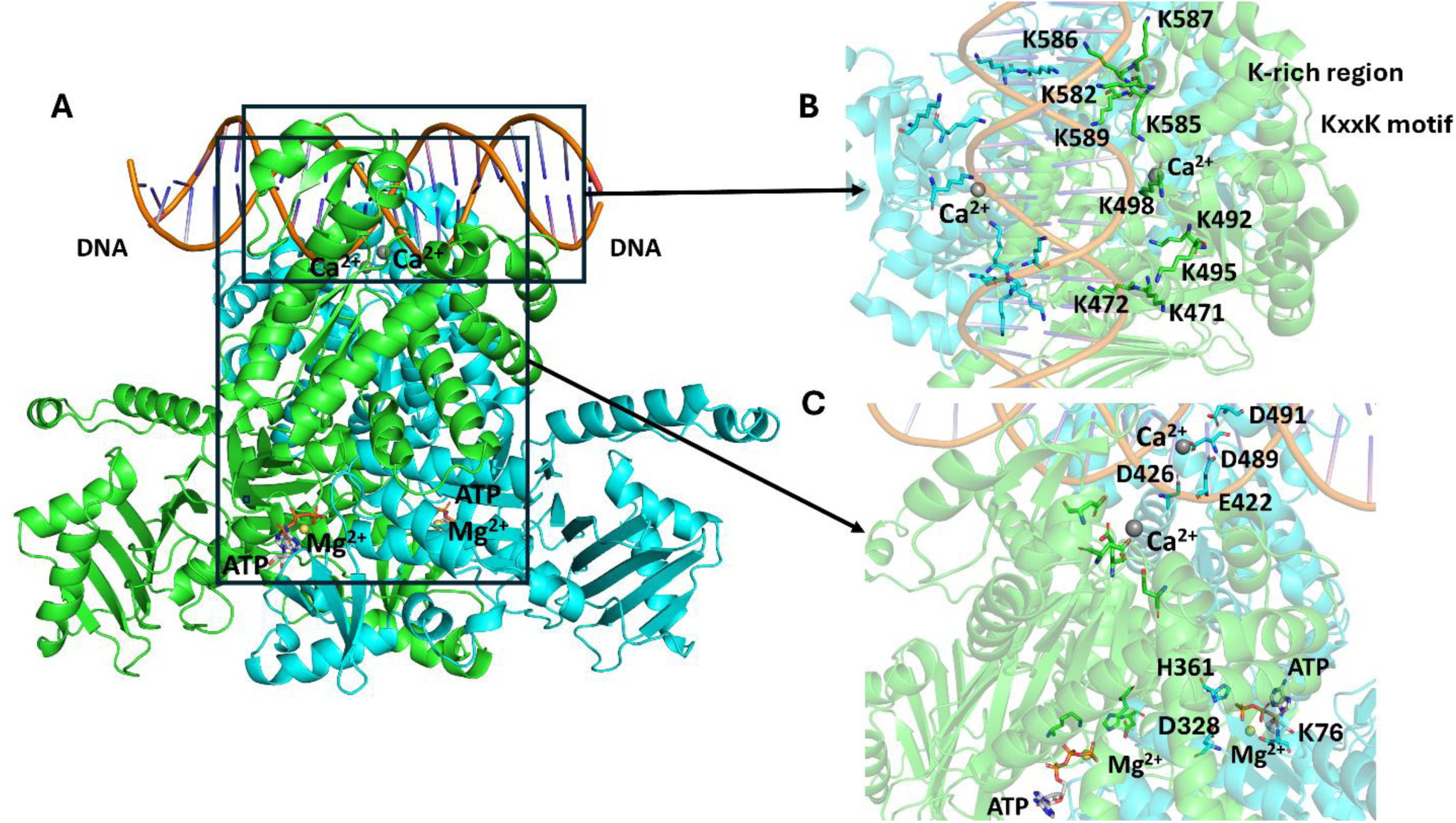
Predicted homodimer of OLD-like protein of bivalve *M. gigas* associated with Ca^2+^, Mg^2+^, ATP and dsDNA. (**A**) Overall structural model of the *M. gigas* OLD-like protein homodimer shown in cartoon representation, with the two monomers colored in green and cyan. The ATP molecules and Mg^2+^ ions (yellow-orange spheres) are located at the central ATPase domains, while the dsDNA is bound at the top Toprim domains and Ca^2+^ ions (gray spheres) coordinate the nuclease active site near the DNA-binding interfaces. (**B**) Detailed view of the DNA-binding interface, highlighting the positively charged K-rich region and KxxK motif. Key lysine residues (including K471, K472, K492, K495, K498, K582, K585, K586, K587, and K589) are depicted as sticks, mediating electrostatic interactions with the negatively charged phosphate backbone of the dsDNA. This architecture is structurally consistent with the DNA-bound state of *B. cereus* GajA (PDB 8X51). (**C**) Close-up views of the catalytic centers. The upper panel shows the Toprim nuclease active site where conserved acidic residues (E422, D426, D489, and D491) coordinate Ca^2+^ ions essential for DNA endonuclease activity. The lower panel displays the nucleotide-binding pocket within the ATPase domain, illustrating the coordination of ATP and Mg^2+^ ions by key residues K76, D328, and H361.

The engagement of the dsDNA is mediated by highly specific electrostatic interactions at the Toprim-DNA interface (**Fig. 4D** and **Fig. 5B**). The negatively charged phosphate backbone of the dsDNA interacts directly with a positively charged, conserved ^492^K-x-x-K^495^ motif and a neighbouring lysine-rich region. This interface is structurally stabilized by key lysine residues, including K471, K472, K498, K582, K585, K586, K587, and K589, which project their sidechains to anchor the nucleic acid. This specific architecture is structurally consistent with the DNA-bound state of the *B. cereus* GajA defense complex assembly (PDB ID: 8X51) (Li et al., 2024).

Ligand binding markedly enhanced the AF3 accuracy of the structural models. Compared to the apo homodimer (pTM: 0.79, ipTM: 0.77), the Mg^2+^, Ca^2+^, ATP-bound homodimer model achieved superior scores (pTM: 0.83, ipTM: 0.80), which were further maximized upon the inclusion of dsDNA (pTM: 0.87, ipTM: 0.86). Furthermore, while the Mg^2+^, Ca^2+^, ATP-bound complex exhibits a minimal structural deviation from the apo state (RMSD of 0.469 Å over 1248 aligned amino acids), the addition of dsDNA induces an extensive conformational realignment, yielding an RMSD of 3.055 Å over 1240 aligned amino acids. This pronounced structural shift suggests that dsDNA engagement drives major conformational rearrangements within the Toprim domains (**Fig. 4** and **Table S4**). These observations align with these of Tang et al. demonstrating that AF3 successfully replicates experimental ligand-dependent conformational transitions driven by nucleotide and substrate binding (Tang et al., 2026).

Ultimately, the predicted dsDNA-bound architecture of the *Mg*OLD-like homodimer shares a closely aligned, overlapping dsDNA binding trajectory within the TOPRIM domains with the experimental cryo-EM structures of the *B. cereus* VD045 GajA defense complex (PDB IDs: 8X51 and 8X5I), despite substantial overall structural divergence (**Fig. S3**). This finding illustrates the profound structural conservation of this immune defense machinery spanning from prokaryotes to certain aquatic eukaryotes (Li et al., 2024).

## 4. Conclusion

This study establishes that the prokaryotic OLD anti-phage defense system is conserved within specific aquatic unicellular lineages and early invertebrates. The highly restricted, mosaic distribution of these elements likely reflects a strict ecological or physiological filter (Ledvina et al., 2024). Genomic analysis revealed that eukaryotic *old-*like genes vary in copy number and intron configurations, while phylogenetic analysis resolves them into three distinct clusters that point to a dual evolutionary trajectory. Cluster 1 likely represents an ancient vertical inheritance derived directly from the ancestral *Promethearchaeota* host during eukaryogenesis (Leão et al., 2024). Conversely, Clusters 2 and 3 indicate independent cross-kingdom horizontal gene transfer (HGT) events. Structural modeling reveals that despite displaying a difference in topology compared to prokaryotic orthologs, eukaryotic OLD-like proteins strictly preserve their fold, catalytic ATPase and Toprim domains, including key catalytic residues. Consequently, these proteins are predicted to be enzymatically active and function as homodimeric to cleave double-stranded DNA. Future functional studies will be essential to validate these predictions and determine whether eukaryotic *old*-like genes retain anti-phage or immune defense capabilities similar to their prokaryotic counterparts.

## Acknowledgments

I thank members of the Bioinformatics Platform at the Institut de Biologie Intégrative et des Systèmes (IBIS), Université Laval, Quebec, Canada for their assistance.

## Data availability

The data that support the findings of this study are available from the corresponding author upon reasonable request.

## Conflict of interest

The author declares no competing interests.

## Funding

No funding

## Supplementary Material

**Table S1**. OLD-like proteins obtained by DELTA-BLAST against the eukaryotic Clustered NR protein database using as query *T. scotoductus* OLD (UniProt ID: E8PLM2).

**Table S2**. OLD-like protein sequences length and taxonomy.

**Table S3**. Determination of the presence of introns and transcripts in *old*-like genes.

**Table S4**. AF3 prediction of *M. gigas* OLD-like homodimers with different combinations of Mg^2+^, Ca^2+^, ATP and dsDNA.

**Fig. S1. Multiple sequence alignment of all OLD-like proteins from aquatic eukaryotes**. Conserved functional motifs of *T. scotoductus* class 1 OLD are indicated by distinct symbols and colors: Walker A (P-loop) (green stars), Q-loop (blue triangles), signature sequence (pink triangles), Walker B (green triangles), D-loop (cyan triangles), H-loop (orange triangles), Metal A and Metal B motifs (blue stars), Metal A motif (D-x-D, green stars), Metal B motif (S-x-E, cyan stars) and spatially conserved basic residue R487 residue (brown star) required for OLD nuclease cleavage. Secondary structures of *T. scotoductus* (PDB ID: 6P74) and *M. gigas* (predicted by AF3) are indicated above and below the MSA, respectively. Accession numbers for all sequences are listed preceding the respective species names.

**Fig. S2. Topology diagrams**. (A) *M. gigas*, (B) *B. cereus* and (C) *T. scotoductus*. β-strands and α-helices are represented as pink arrows and red cylinders, respectively. N-terminal and C-terminal of sequences are represented by N and C letters in yellow, respectively.

**Fig. S3. Predicted homodimer models of the *M. gigas* OLD-like protein bound to Ca²⁺, Mg²⁺, ATP, and dsDNA.** (**A**) Front and (**B**) top views of the *M. gigas* homodimer complex. Monomer A is shown in green, monomer B in cyan, dsDNA in orange, and ATP in light gray. Ca²⁺ and Mg²⁺ ions are represented as gray and yellow-orange spheres, respectively. (**C**, **D**) Structural superposition of the *M. gigas* homodimer-dsDNA with the *B. cereus* GajA homodimer (in pink) in complex with dsDNA and Ca²⁺ (PDB ID: 8X51). Note the close spatial alignment and overlapping paths of the bound dsDNA molecules.

## Notes

### Competing Interest Statement

The authors have declared no competing interest.

### Summary of Updates

Nothing has been changed in the article except for the addition of the concluding paragraph..

