## Supplemental tables_Figs for "Certain Aquatic Eukaryotes Harbor an OLD-Like Immune Defense System": Figure S1_MSA_OLD-like_proteins.pdf

```
E8PLM2_Thermus_scotoductus                                                                   β1
                                                                                   1          10
E8PLM2_Thermus_scotoductus                            ......................................MLKRL.......QVKNFRCL.
UJG41740.1_Candidatus_Heimdallarchaeum_aukensis       .....................................MILKKI.......YIKNFLSYS
MFW9996705_Candidatus_Odinarchaeota_archaeon          .....................................MYLSKI.......YLENFLSYE
MCH8915407.1_Thaumarchaeota_archaeon                  .....................................MKIDWI.......KISNVFSFK
MHA1972560.1_Candidatus_Hodarchaeales_archaeon        M....................................AKIIKF.......RIKNYKSIK
NHI95076.1_Candidatus_Lokiarchaeota_archaeon          MDTTD....................LTQLPLTTPSHNLKLKKI.......KYENFKTF.
MEM2144305.1_Candidatus_Jordarchaeaceae_archaeon      .....................................MPVNRF.......EIKNTGPI.
AHB41284.1_candidate_division_SR1_bacterium           .....................................MQIISA.......NISNILSFP
HEV2339438.1_Patescibacteria_group_bacterium          .....................................MKINYI.......QISNILSFE
MBL9158386.1_Verrucomicrobiales_bacterium             M..................................PTIMLKTL.......TIQNFKRFK
MGB8583703.1_Candidatus_Sulfotelmatobacter_sp.        .....................................MKLLGA.......EFSDFACL.
MBI3627515.1_Candidatus_Sungbacteria_bacterium        .....................................MKLKKF.......KIKNYKSIV
_213105201.1_Candidatus_Protochlamydia_amoebophila    M..........................................................
CDRQKP010002909.1_Symbiodinium_sp.                    MKKLL.........NNWTLGKIRFK.........MEDIEILSY.......RIKNYKSIK
JAUKPS010110286.1_Chlorellidium_tetrabotrys           .....................................MRVVKF.......RVWNYKSII
XP_004364654.1_Capsaspora_owczarzaki                  MVELD.........EPLSLLEELFDRDDPPAKQDKPNCFLLSL.......RLQNFKRF.
KJE91799.1_Capsaspora_owczarzaki                      MT...................................GFLTSL.......QLTNFRKF.
XP_048575736.1_Nematostella_vectensis                 ......................................MLRRV.......VVENYVIFK
A0A6P8J3J2_Actinia_tenebrosa                          ......................................MLRRV.......VVENYVIFK
XP_022780642.1_Stylophora_pistillata                  ......................................MLRRV.......VVEDYIFFK
A0A9X0CFV3_Desmophyllum_pertusum                      ......................................MLRRI.......VVENYIIFK
A0A6S7HLM0_Paramuricea_clavata                        MM.................................AGATLKRV.......AIDNYKVFN
XP_046856289.1_Xenia_sp.                              M....................................ATLTRV.......LVENYKVFC
A0A813MIX4_Adineta_steineri                           M...................................GSKLKSLVSVGEVPSLVQYKEFH
UJR19038.1_Adineta_vaga                               MARCDSTTCDKMNSSPEDIDQSRHLTLENGIIRKPVGMILKKI.......HIFKYRQFE
JAXIUW010014154.1_Cephalothrix_simula                 ......................................MLRSV.......VFSNFIHFK
A0A8B8DK01_Crassostrea_virginica                      ......................................MLKRI.......VFENFVHFS
XP_070564142.1_Ptychodera_flava                       ......................................MLRRV.......IVENYVNFT
ACQM01009845.1_Saccoglossus_kowalevskii               MIYII.........................FFVVVPVNMLSRV.......AIDNYVNFN
XP_078582628.1_Branchiostoma_floridae_japonicum       MASEE..............................QGSVLTEI.......SFKNFCRF.
JALCYT010000124.1_Branchiostoma_belcheri              ............................NFFFHLPTGSLLTEI.......SFENFCRF.
KAJ3014353.1_Thoreauomyces_humboldtii                 .....................................MHIVSL.......NLDSLCKF.
KAJ3188706.1_Gaertneriomyces_sp._JEL0708              M....................................VYIQTL.......HLLAFRRF.
XP_002110806.1_Trichoplax_adhaerens                   MASFQR....PHNRNETEMYQEHY..DNEIDKKCKKLGYLTEIK.....ITGNGFKNLD
RDD37009.1_Trichoplax_sp._H2                          MASFQR....PHNRNETEMYQERY..DNEIDKKCTMAGYLSEMK.....VTGNGFKNLN
A0A8W8L770_Magallana_gigas                            ......................................MLKSI.......TLENFVHFK
A0A8W8L770_Magallana_gigas                                                                TT
                                                                                   β1

                                                            β2                                                  β3
E8PLM2_Thermus_scotoductus                            ......
                                                                20
E8PLM2_Thermus_scotoductus                            ......EDIDLPLG........................................PLTAI
UJG41740.1_Candidatus_Heimdallarchaeum_aukensis       ......EEQEINLN........................................QKSII
MFW9996705_Candidatus_Odinarchaeota_archaeon          ......KAVSFDVE........................................SKLVI
MCH8915407.1_Thaumarchaeota_archaeon                  YHENIDQGPTIVFD.......................................KNLNIL
MHA1972560.1_Candidatus_Hodarchaeales_archaeon        ......DSGDCYLS.......................................NNITVL
NHI95076.1_Candidatus_Lokiarchaeota_archaeon          ......QKLEVELS........................................DFNIL
MEM2144305.1_Candidatus_Jordarchaeaceae_archaeon      ......KYARIDNL.......................................PSVVII
AHB41284.1_candidate_division_SR1_bacterium           YQEQFSEERKIHFD.........................V..........NKKGGLNIL
HEV2339438.1_Patescibacteria_group_bacterium          YRDDINHSPKINFD.......................................KGLNIL
MBL9158386.1_Verrucomicrobiales_bacterium             ......QRTKIDFE........................................AITIL
MGB8583703.1_Candidatus_Sulfotelmatobacter_sp.        ......DRQFIPLR.......................................PGINLL
MBI3627515.1_Candidatus_Sungbacteria_bacterium        ......DSGDCYLT.......................................DTITIL
_213105201.1_Candidatus_Protochlamydia_amoebophila    .......................................................FSVI
CDRQKP010002909.1_Symbiodinium_sp.                    ......DSGEIYIK.......................................TPITIF
JAUKPS010110286.1_Chlorellidium_tetrabotrys           ......DSGDCFPA.......................................EKVTIL
XP_004364654.1_Capsaspora_owczarzaki                  ......HDLVFKFS.......................................SSPKVI
KJE91799.1_Capsaspora_owczarzaki                      ......KDETFHLT.......................................TSPKVI
XP_048575736.1_Nematostella_vectensis                 ......GRHELDFM.....................GRD..............GCNFYTF
A0A6P8J3J2_Actinia_tenebrosa                          ......GRHELDFD.....................GKE..............GRHFYTF
XP_022780642.1_Stylophora_pistillata                  ......GRQELNLE.....................SSS..............GNSFHTL
A0A9X0CFV3_Desmophyllum_pertusum                      ......GRQELDLD.....................STC..............GGSFHTL
A0A6S7HLM0_Paramuricea_clavata                        ......NKQVLDFRHGP.................SLST..............NSKYFTL
XP_046856289.1_Xenia_sp.                              ......TKQEVDFRQEL.................ASST..............SNKYFTL
A0A813MIX4_Adineta_steineri                           SNGN..QVPELNLA.......................................KRITVL
UJR19038.1_Adineta_vaga                               SAEQ..GEYVFHLA.......................................DHVTAL
JAXIUW010014154.1_Cephalothrix_simula                 ......ERQYVRFQ.......................................NGPNYI
A0A8B8DK01_Crassostrea_virginica                      ......EKNIIDFT....................SSEN..............NAALNIF
XP_070564142.1_Ptychodera_flava                       ......ELQDLNFS.....................PKH..............QHNLYLF
ACQM01009845.1_Saccoglossus_kowalevskii               ......NPQDLKFC....................L..Q..............QDGIHLF
XP_078582628.1_Branchiostoma_floridae_japonicum       ......ENVTIDLH.......................................SESISL
JALCYT010000124.1_Branchiostoma_belcheri              ......KKVTINLD.......................................SESTSL
KAJ3014353.1_Thoreauomyces_humboldtii                 ......DRTDIPLS........................................ELTVL
KAJ3188706.1_Gaertneriomyces_sp._JEL0708              ......SDLHIELA.......................................GSLATF
XP_002110806.1_Trichoplax_adhaerens                   ...............................QSFCWSNI..............PMLAVI
RDD37009.1_Trichoplax_sp._H2                          ...............................QSFCWSNI..............PMLAVI
A0A8W8L770_Magallana_gigas                            ......DRTVIDFDTTPRVKQSKASLHKEATSSNTSENKKQRRDAEKEVASGWNTLNIF
A0A8W8L770_Magallana_gigas
                                                               β2                                                β3
```

```
                                                   α1               η1         η2
E8PLM2_Thermus_scotoductus                         β1 (►)       ℓℓℓℓℓℓℓℓℓℓℓℓ         ℓℓℓ        ℓℓℓ                  ▬..
                                                      30         40         50         60                    70

E8PLM2_Thermus_scotoductus                         VGPNGAGKTTILRAIDLVLGDVWPSLRSFRIPQDFINF.............DTTRAI..
UJG41740.1_Candidatus_Heimdallarchaeum_aukensis    VGENGSGKSNLVRIIDFILSNSATSNLDWSYFPFW................DTKTQS..
MFW9996705_Candidatus_Odinarchaeota_archaeon       VGANGSGKSNILRFFRTLLGVEPNYNLYWKDHKSW................DKTKPC..
MCH8915407.1_Thaumarchaeota_archaeon               IGPNGSGKSNFLEIINQVFRKFFFRDCPLSLEKTIEL.......QSLS..GNDRTNQLN
MHA1972560.1_Candidatus_Hodarchaeales_archaeon     AGRNESGKTSILEALEDFDTEKAIRQDAVPIH...................DSNLKP..
NHI95076.1_Candidatus_Lokiarchaeota_archaeon       VGTNNSGKTNILLGIYSFFEFLKQLYSQHEDLKVN................DS......
MEM2144305.1_Candidatus_Jordarchaeaceae_archaeon   AGPNGVGKSTLLETLKKQSGIFMEGTGKLLYIPPY........................
AHB41284.1_candidate_division_SR1_bacterium        IGPNGSGKTNFLEIITQLIKVGLIKDFVYTENNGIQNSI..........IENQWPLE..
HEV2339438.1_Patescibacteria_group_bacterium       IGPNGVGKSNFLEILNQLFKNILYRQAIYSEDNVF................NHRSNG..
MBL9158386.1_Verrucomicrobiales_bacterium          LGANNSGKSTVLQALSIFQYCLDNTRKKKNGALVL................DTKTIG..
MGB8583703.1_Candidatus_Sulfotelmatobacter_sp.     VGKNNAGKTAILRGLSALAALPVGTVQPMTTE.................LSGYCRDA..
MBI3627515.1_Candidatus_Sungbacteria_bacterium     AGKNESGKTSILEALQDFDTDEKIRESAKPIK...................SPEAVP..
_213105201.1_Candidatus_Protochlamydia_amoebophila AGKNGVGKSNLLLAILFGSFSNQGTIKLSSNNFQICSLKNTDELKTLRQ..DTENKS..
CDRQKP010002909.1_Symbiodinium_sp.                 AGMNESGKTNFLEAFYDINVEREINNLSKRIN...................DDES....
JAUKPS010110286.1_Chlorellidium_tetrabotrys        AGKNEAGKSSLLEALADFNPDLPIRENAVPLEKQ.................RSNQKP..
XP_004364654.1_Capsaspora_owczarzaki               AGRNGFGKTQLLWAIMVYLRAYNARVEKNSTWHS.................NGERYL..
KJE91799.1_Capsaspora_owczarzaki                   MGRNGSGKTQLLWAIAIFLRSYNTRVATSNWH...................KQNKPTL.
XP_048575736.1_Nematostella_vectensis              VGENASGKSSLVALIKAASNLQEEGPDFEVI....................DNCKES..
A0A6P8J3J2_Actinia_tenebrosa                       VGENASGKSSLVALIKAACNLQDEGPDFEVIT...................NGKGES..
XP_022780642.1_Stylophora_pistillata               VGENATGKSSFVALVKATSNFDNNENDFEVIA...................NKETGS..
A0A9X0CFV3_Desmophyllum_pertusum                   VGENASGKSSFVALVKAASNFVDNDEDFEVIG...................NDIASA..
A0A6S7HLM0_Paramuricea_clavata                     VGENGSGKSTFLALVGLASDFAGKVTNH.....................VCDQAKGS..
XP_046856289.1_Xenia_sp.                           VGENGSGKSTFLALIGLASNFGGKVTSH.....................ACDKTKAS..
A0A813MIX4_Adineta_steineri                        VVPNSCGKTTILQAVYLFCHAVRRGSGEYSFK..........EMTTLFWAKNALSYG..
UJR19038.1_Adineta_vaga                            VGPNSSGKTTLLKMMRLFCAAVKFGSTCLPIAYAE......KELGIDEEWLNGDFHV..
JAXIUW010014154.1_Cephalothrix_simula              VGGNSTGKSSIFELIRRCLSDTISKSYST....................ISSKDQAA..
A0A8B8DK01_Crassostrea_virginica                   VGANHSGKSTVCELIRRCMTNEINVTQTRPF....................NKDYVA..
XP_070564142.1_Ptychodera_flava                    VGENGSGKSSLLELIEYAALDFSSKEITEKIN.................VRDKKKDA..
ACQM01009845.1_Saccoglossus_kowalevskii            VGENGSGKSSLVELIEYASLDYSPRKRIKNIN.................VCNRQRDA..
XP_078582628.1_Branchiostoma_floridae_japonicum    LGENGRGKTTVLKGIELFANLFDSYMTDEENVPSRSS.......NLVLI..NEKKFL..
JALCYT010000124.1_Branchiostoma_belcheri           LGENGRGKTTVLKGIELFATLFDSFANSTERTPYS...........VTY..QKQDFS..
KAJ3014353.1_Thoreauomyces_humboldtii              VGTNRSGKTSVLQGIGLLALLMRNTENDGTITAGHGG............GSDYIKRMAD
KAJ3188706.1_Gaertneriomyces_sp._JEL0708           VGTNGSGKTSLLQAVAVFSYLVEYATAVDGTLQSS..........TLK.IHDDARQVLD
XP_002110806.1_Trichoplax_adhaerens                TGKNGCGKTAVLEAIMRGFREEKENQSSKNS............................
RDD37009.1_Trichoplax_sp._H2                       TGKNGCGKTAVLEAIMRGLREEEARRLTEGSK...........................
A0A8W8L770_Magallana_gigas                         VGANFCGKSTVIELIRRCMTDEINVSVTNSF....................DDNSFA..
A0A8W8L770_Magallana_gigas                         ──► TT  ℓℓℓℓℓℓℓℓℓℓℓℓ                                  TT  ▬..
                                                               α2
                                                     ★★★★★★★★

                                                      β4                β5                  β6
E8PLM2_Thermus_scotoductus                         ────────────►TT──────────────►T....T──────────►TT        TT
                                                            80         90         100        110        120

E8PLM2_Thermus_scotoductus                         EITVHFDPPYTQGSFNITAFRLTCKGED...ADFHVDLEPLDEGGNVPRYPSGNP...
UJG41740.1_Candidatus_Heimdallarchaeum_aukensis    KIYLTFELSEDERIFLRKFLVLSFCSMH.........LSSDQEFGRALIYSFFNNWEN
MFW9996705_Candidatus_Odinarchaeota_archaeon       KCYCRFKLTTEDKEFFKYCLVLQLISMDNSD......REEREEIGNYLLKELTNP...
MCH8915407.1_Thaumarchaeota_archaeon               NITLQ.....KNHNF.......................QKNNNFKEKPFEI.......
MHA1972560.1_Candidatus_Hodarchaeales_archaeon     EISITFELEKEEFGKISENIIAD..............KEN..................
NHI95076.1_Candidatus_Lokiarchaeota_archaeon       KANCEYFFLKVPEDAQI.........................................
MEM2144305.1_Candidatus_Jordarchaeaceae_archaeon   RTPVTFSLHKS...............................................
AHB41284.1_candidate_division_SR1_bacterium        NMIPHFSYQDKPSIV...........................................
HEV2339438.1_Patescibacteria_group_bacterium       QVPLGQTITEAPRNYNY....................LNPNRKSTNSVRKI.......
MBL9158386.1_Verrucomicrobiales_bacterium          .........................PD..EFGALPV.ASPKDLWSN............
MGB8583703.1_Candidatus_Sulfotelmatobacter_sp.     KSTFTFELIYRTEERDADFF...........HHPAL......................
MBI3627515.1_Candidatus_Sungbacteria_bacterium     EISITFVVEKETLKEILEEI.................KSPVKATA.............
_213105201.1_Candidatus_Protochlamydia_amoebophila KVKLQAELEKSFQDLNEY........................................
CDRQKP010002909.1_Symbiodinium_sp.                 EVNLILNIKIKDFNI......................IEEYDLDGDKIIFIKTKE.GY
JAUKPS010110286.1_Chlorellidium_tetrabotrys        KIEIVFESTAEALIKIIDKLGLDLPGRQ..SF.AASIGKSP.................
XP_004364654.1_Capsaspora_owczarzaki               PLTSLFGDPYFEQQD...........................................
KJE91799.1_Capsaspora_owczarzaki                   ALGPLFGDPFFEYET...........................................
XP_048575736.1_Nematostella_vectensis              KVVCEFEFKNGAELIRH....................CDH..................
A0A6P8J3J2_Actinia_tenebrosa                       KAVCHFEFSNGKELVKV....................CEES.................
XP_022780642.1_Stylophora_pistillata               KAVCEFHFENGEELSGY....................VDKMH................
A0A9X0CFV3_Desmophyllum_pertusum                   KAVCEFHFKHGHELLQY....................FERS.................
A0A6S7HLM0_Paramuricea_clavata                     VISCEYVIENWLELT...........................................
XP_046856289.1_Xenia_sp.                           MVACEYKVENWAEIM...........................................
A0A813MIX4_Adineta_steineri                        RLSGTFEVFYKD..................I......LETKELKLDVRFFQ.......
UJR19038.1_Adineta_vaga                            RLQGTFVVCYENDNQ...............V......IPCSDLTLNVSLRVSDTC...
JAXIUW010014154.1_Cephalothrix_simula              YAICEFELPDGQFQY..................SGVLVEKDDNQTRNMFKLSMKA...
A0A8B8DK01_Crassostrea_virginica                   YAFCDFRLDAKEEVICI................SGMIKEPDEKDAYKIFWFKDKS...
XP_070564142.1_Ptychodera_flava                    LMLCSFEIDTEDEDMLTSELHRCL.............RDNHKDCGD............
ACQM01009845.1_Saccoglossus_kowalevskii            YMLCDYDISQKEWRKLYEHCQRVDT.......HGDV.DEDDGNDGDNASDHEESK...
XP_078582628.1_Branchiostoma_floridae_japonicum    ...............................SWLGV.TELKLIWYN............
JALCYT010000124.1_Branchiostoma_belcheri           ...............................NLLGI.TEPDLLWHK............
KAJ3014353.1_Thoreauomyces_humboldtii              QVAL......................................................
KAJ3188706.1_Gaertneriomyces_sp._JEL0708           TLSMS.....................................................
XP_002110806.1_Trichoplax_adhaerens                ..........................................................
RDD37009.1_Trichoplax_sp._H2                       ELKIEH....................................................
A0A8W8L770_Magallana_gigas                         YVFCQFNLSPYDEVI..................SGIIKEPGNDVMYKVFIYSAKN...
A0A8W8L770_Magallana_gigas                         ─────────►TT─────.....................────►  TT  ─────────►TT
                                                       β4                  β5                     β6
```

*E8PLM2_Thermus_scotoductus*

```
E8PLM2_Thermus_scotoductus                          ...........................L...............................
UJG41740.1_Candidatus_Heimdallarchaeum_aukensis     FLKDSFSDLTIGFCSESNNPER.....L.............KYFLILSFNESLNI....
MFW9996705_Candidatus_Odinarchaeota_archaeon        ...SFMEYIDVFLSSNTTDPEE.....L.............KTSIIFPVNTNLHAC...
MCH8915407.1_Thaumarchaeota_archaeon                .......KIQIKFSPKDRENFQ.....F...............................
MHA1972560.1_Candidatus_Hodarchaeales_archaeon      .........RIILKKSFDNEYS.....IINTEELF.......................S
NHI95076.1_Candidatus_Lokiarchaeota_archaeon        .......WYNKKWMSRKYIPVK.....F...EFEFTDG.....................
MEM2144305.1_Candidatus_Jordarchaeaceae_archaeon    ...........................L...............................
AHB41284.1_candidate_division_SR1_bacterium         .......DMEFHVSENDKENML.....F...............................
HEV2339438.1_Patescibacteria_group_bacterium        .......KISLNLNQNDKNNLK.....F...............................
MBL9158386.1_Verrucomicrobiales_bacterium           ..............GQSSKPIE.....I...EAEFDNG.....................
MGB8583703.1_Candidatus_Sulfotelmatobacter_sp.      .......SATDPLMQDTVQKML.....L...RELLSEETTAR.................
MBI3627515.1_Candidatus_Sungbacteria_bacterium      .......DVEFEIIKTFPDKYTFGETTL..QSEIFGGKEIEKI...............Q
_213105201.1_Candidatus_Protochlamydia_amoebophila  .......AVNKLLRKGSDIPSA....................................Q
CDRQKP010002909.1_Symbiodinium_sp.                  ....EIENRTKLLNKKYLINRK.....V...............................
JAUKPS010110286.1_Chlorellidium_tetrabotrys         .........ELTITKSYPQTYE.....V..SAGIFSEI..QKIKPSD..........WS
XP_004364654.1_Capsaspora_owczarzaki                .G.....TVYKSLITSARDTAT.....L...TGTFGPKS....................
KJE91799.1_Capsaspora_owczarzaki                    .......DTSSALIRQGETTAT.....L...LGNFSDG.....................
XP_048575736.1_Nematostella_vectensis               .............CNEASATQL.VSPGL...SSWFSDDILK.......IPTPFFDQIFE
A0A6P8J3J2_Actinia_tenebrosa                        .............CATAGQATK.LSVGL...SSWFCDELIK.......IPSALLEPIKM
XP_022780642.1_Stylophora_pistillata                ...........SYSAKALLPNFPFISGL...FSWFSDYILI.......FPVQLINLLQE
A0A9X0CFV3_Desmophyllum_pertusum                    .............STRALHPTVPCVSGL...FSWFSDDILK.......FPMQLADLLRD
A0A6S7HLM0_Paramuricea_clavata                      ..............NTHSNVVH.NYPVL...ELWFLTGVLK.......IPHMLQQYISE
XP_046856289.1_Xenia_sp.                            ..............STSKVPGQ.SYPAL...QIWFMGGLLK.......IPPDLHGFLTE
A0A813MIX4_Adineta_steineri                         .......NRTSHEDKNHNEHKP........YSYVFVEAP....................
UJR19038.1_Adineta_vaga                             ......KGYSVYIDVNNSINAA.....I...NAAYRKAGKNKSE...............
JAXIUW010014154.1_Cephalothrix_simula               ....DKCDVDVYDYSDVRDETG........RCAPYEKS..............LHIT.KT
A0A8B8DK01_Crassostrea_virginica                    .......GIYLRLKCSGKNEFT........KKNPFEDIEQKF.................
XP_070564142.1_Ptychodera_flava                     .....GEEEEEAGEKQEEGNKK.....L..FAGFFFDHMTGDTV.............KF
ACQM01009845.1_Saccoglossus_kowalevskii             ....DGKSVGNANQSHKKDGISYN...L..CTGFIYQSSTNKSG.............KF
XP_078582628.1_Branchiostoma_floridae_japonicum     ..............TDISKSIN.....I...CGKLSD......................
JALCYT010000124.1_Branchiostoma_belcheri            ..............RDTNESIK.....I...SGKLSYKGKSLD................
KAJ3014353.1_Thoreauomyces_humboldtii               .......GNSDYLSRQGDVPVS.....I...GATFSDG.....................
KAJ3188706.1_Gaertneriomyces_sp._JEL0708            .......GDRSLLCRNGEEPVH.....L...GATLTNE.....................
XP_002110806.1_Trichoplax_adhaerens                 .......KISSKEGYRSFKGVT.....I...............................
RDD37009.1_Trichoplax_sp._H2                        .......KKSIKHNGRTDKDIA.....I...............................
A0A8W8L770_Magallana_gigas                          .......DTFLR.SKSSESIAT........YSGLVHEADDKE.................
A0A8W8L770_Magallana_gigas                                 ━━━━━━━.━━➤TT━━━━━━.......━━━➤   ℓℓℓℓℓ.............
```

*E8PLM2_Thermus_scotoductus*

```
                                                                                                        α2
                                                                                                     ℓℓℓℓℓℓ
                                                                                                      130
E8PLM2_Thermus_scotoductus                          ................................................RVGTDMRNHAR
UJG41740.1_Candidatus_Heimdallarchaeum_aukensis     .YLSNPSTNIIIILVDKEKK............................LDEILTLSKNQ
MFW9996705_Candidatus_Odinarchaeota_archaeon        .YLITRNNELLLLKDYQEG............................NIEAIEAAQITV
MCH8915407.1_Thaumarchaeota_archaeon                .IFQHRDKIVGMLKEYYEIGTSFQISTITEQDLTNLENLEFSISE.....SIDPRQNRR
MHA1972560.1_Candidatus_Hodarchaeales_archaeon      KIIKPNKSRIKLFEKNRTD..........LKKIFDRNNIKLEL.........DAINEGH
NHI95076.1_Candidatus_Lokiarchaeota_archaeon        ....................................LLIELHLYRYFGVPSIKLINLNK
MEM2144305.1_Candidatus_Jordarchaeaceae_archaeon    PFIGPRTRFVDILTSDNFSI...........HAPGISLPHYIISGG.............
AHB41284.1_candidate_division_SR1_bacterium         .IQKKQEIFSKIIETYSTTK..YKIPVCDIEKIKNLQTLHIQFT.......IDTTNKTA
HEV2339438.1_Patescibacteria_group_bacterium        .IFDNATDFNTIFSNYSNVGGLSFETNFNKQDIDNTNDIEFEYE......DTNPINQPA
MBL9158386.1_Verrucomicrobiales_bacterium           ....................................SKIKFEIKFQYNRFSIHPTVTGP
MGB8583703.1_Candidatus_Sulfotelmatobacter_sp.      ......................WRFKVFGMNRLVVFLDLTFEFT......PSEKESRKA
MBI3627515.1_Candidatus_Sungbacteria_bacterium      KSIKDKWEKIKLIRGKYEA..........LAG....NLFDFEFS.......NFENDKTL
_213105201.1_Candidatus_Protochlamydia_amoebophila  PLFDEVIESTLQVINKKDSN..............SFSNFEHELKNQ.............
CDRQKP010002909.1_Symbiodinium_sp.                  .....IEDNLKTIKNHFPS..........IKD....NEIFKEI.........EKENNKT
JAUKPS010110286.1_Chlorellidium_tetrabotrys         TIFAALSAVVSLAQTAAAA..........KKLGL..PKLGYEIY.......TEKDSAAA
XP_004364654.1_Capsaspora_owczarzaki                ...............................................WPTEIQVQLRAT
KJE91799.1_Capsaspora_owczarzaki                    ....................................RKVSFVIN.......PAKSNRAR
XP_048575736.1_Nematostella_vectensis               .YCRGGTKRIQVLAGHRHQS................LPQTTESSRR.KENEVDRISNSQ
A0A6P8J3J2_Actinia_tenebrosa                        .ACRHVNSKLDLLVGHRHQ..................PIGTQYSRPAYDPELDPLCNSQ
XP_022780642.1_Stylophora_pistillata                .TFGSASSLIRVFCGRRNL...............................NTSNPFHSQ
A0A9X0CFV3_Desmophyllum_pertusum                    .TFGSVSALVRVFSGRKSF...............................NNNHPSHCQ
A0A6S7HLM0_Paramuricea_clavata                      KYLASNTKDLTVVIGFKHLP...........................SSESSGEAVVSK
XP_046856289.1_Xenia_sp.                            RYVGLNVKDVLLVIGFRRSA...........................YSASNFEENVHR
A0A813MIX4_Adineta_steineri                         ..................................................NVKQQNTPP
UJR19038.1_Adineta_vaga                             ...............................................EILSTGQSETRE
JAXIUW010014154.1_Cephalothrix_simula               SLLNDLMEHIQIIIKYSDEI................EKEAFKLSDC.NFQEIAEKEKSI
A0A8B8DK01_Crassostrea_virginica                    ...............................................DISIFDKKEKVE
XP_070564142.1_Ptychodera_flava                     YFLKSQTSCIACIEGYGDI.................DNISHEDK...............
ACQM01009845.1_Saccoglossus_kowalevskii             SILIAGTNAVTYV.......................................EVPQNAS
XP_078582628.1_Branchiostoma_floridae_japonicum     ....................................KRIEFEVS......KREQGTQQV
JALCYT010000124.1_Branchiostoma_belcheri            ....................................KPIQFEVS.......RQEGT..V
KAJ3014353.1_Thoreauomyces_humboldtii               ....................................SSVKYAFETT.............
KAJ3188706.1_Gaertneriomyces_sp._JEL0708            ....................................VKIMFRLSDV.............
XP_002110806.1_Trichoplax_adhaerens                 ...........................................................
RDD37009.1_Trichoplax_sp._H2                        ...........................................................
A0A8W8L770_Magallana_gigas                          ...............................................AIHSIVEKESVN
A0A8W8L770_Magallana_gigas                          ...............................................ℓℓℓℓℓℓ
```

```
                                                    β7                               β8         α3
E8PLM2_Thermus_scotoductus                          ▬..➔ T..........T                ➔ ℓℓℓℓℓ..ℓℓℓ
                                                         140                      150         160

E8PLM2_Thermus_scotoductus                          VL..FLDH..........RR..................NLAQHLPSIRGSILGR..LLQ
UJG41740.1_Candidatus_Heimdallarchaeum_aukensis     YL..DFRN.......................ITSFVK.SFNDLYPGEDIKELVKN.NED
MFW9996705_Candidatus_Odinarchaeota_archaeon        II..DFSQ.......................MTRFREF....PLPGGL.........DK
MCH8915407.1_Thaumarchaeota_archaeon                QL..KISQ.SLTEIEQFVYDYLVWFEFIQN.IILIANE..............KLSMDWE
MHA1972560.1_Candidatus_Hodarchaeales_archaeon      FS..KISN.QLSILNTDITRLSRLSELDSVTALELIS.TMEQIIGEFV........EPK
NHI95076.1_Candidatus_Lokiarchaeota_archaeon        TL..DKDY.......................IRKILN.SSPIMIPSFIGLLINEEYSTR
MEM2144305.1_Candidatus_Jordarchaeaceae_archaeon    ......................................PRSRSAPDFA........PYF
AHB41284.1_candidate_division_SR1_bacterium         HI..SNKS..KNKIENFAIEYLIYQELFQI.AIMIYNNNIKKS..........DELGRY
HEV2339438.1_Patescibacteria_group_bacterium        FH..RVNR.QQNQCVNFIEKYFEYFEFLQK.AIQIGNT..............FENKQWN
MBL9158386.1_Verrucomicrobiales_bacterium           VS..QLVE.............................TTKIRYVPIHSGLGL...REEY
MGB8583703.1_Candidatus_Sulfotelmatobacter_sp.      LI..LTKN.ANGVLL...LRRVRYPDLVSDASKQVGKS..TIAAPDGAYPVLEPD.QLS
MBI3627515.1_Candidatus_Sungbacteria_bacterium      FT..NFTS.ATTPNV......TQITDEKERERFSKLLDEIQTDINNLA........NLL
_213105201.1_Candidatus_Protochlamydia_amoebophila  ......................................FNKRNTPHIA........DYN
CDRQKP010002909.1_Symbiodinium_sp.                  FY..RNIE.NIEEK.........LISLKSVHNINIEY.DFDKKVDINY........ILE
JAUKPS010110286.1_Chlorellidium_tetrabotrys         VA..EYET.KISELI......GALSEVDQNTIRSNIA.TLKSSLDENS.......RQKI
XP_004364654.1_Capsaspora_owczarzaki                ST..EVRH.DDGQ...........................................VHQ
KJE91799.1_Capsaspora_owczarzaki                    MI..EQPD.TAEQ...........................................PPR
XP_048575736.1_Nematostella_vectensis               FL..YILM.DDGILV........ATKSGGHLITGFHE.....PLPAGV.NPA....DWV
A0A6P8J3J2_Actinia_tenebrosa                        FL..YIIM.DDGIFV........ATKSEGIITTGFHE.....LLPEGV.NPM....DWI
XP_022780642.1_Stylophora_pistillata                FL..YFVS.EEIVLV........VTKDNGILSAALHD.....PLPVGV.DLI....EWS
A0A9X0CFV3_Desmophyllum_pertusum                    FL..YVVS.EDVVLV........VIKNNGELSAALHD.....PLPAGV.DLI....EWC
A0A6S7HLM0_Paramuricea_clavata                      FL..YVQS.GSSIIA.......LVHELNGEAKVAVVE.......GLENTDLL....HWM
XP_046856289.1_Xenia_sp.                            FL..YIQF.KNHVLA.......LIHGLDSEAKILVNE.......PSDVVNLL....HWR
A0A813MIX4_Adineta_steineri                         HI..YLKS...................................................
UJR19038.1_Adineta_vaga                             HI..HIRV.................................NVQPKDT.....HSIQWE
JAXIUW010014154.1_Cephalothrix_simula               TE..RITK.SIDILYDI.....................LEKDSLPFKT.DVTERNISLR
A0A8B8DK01_Crassostrea_virginica                    LFLKKISK.......................LIIIGE.PSDEGVPEKP........DWK
XP_070564142.1_Ptychodera_flava                     ......................................TSHERYPS..........TGS
ACQM01009845.1_Saccoglossus_kowalevskii             FI..ETPT.RATQA..........................HPLLPAVL.....HNKTAT
XP_078582628.1_Branchiostoma_floridae_japonicum     RV..RCADWSRSDPKQE........KIEGPTVVSISN.TGVFTAPFKA...........
JALCYT010000124.1_Branchiostoma_belcheri            KV..KCSH.DRSELK..........EIGCPTVVSISN.TGAFMAPFKT...........
KAJ3014353.1_Thoreauomyces_humboldtii               .........................RLDSGFVVSCAP.AESHKYPPVRYTFFPRDVAYS
KAJ3188706.1_Gaertneriomyces_sp._JEL0708            ..........................HGTDLVVSVEPDVPASSIPRVRCTLFPRDLRFD
XP_002110806.1_Trichoplax_adhaerens                 ......................................KYATEVPKCH...........
RDD37009.1_Trichoplax_sp._H2                        ......................................KCSDEIPKCH...........
A0A8W8L770_Magallana_gigas                          II..QLLH.......................LIKSSN.TQKDCLPEEP........SWK
A0A8W8L770_Magallana_gigas                          ℓℓ..ℓℓℓℓ.......................ℓℓℓℓℓ                     ℓℓ
                                                                      α4
                                                         ▲▲▲          ▲▲                    ▲

E8PLM2_Thermus_scotoductus                          ℓℓℓℓℓ

E8PLM2_Thermus_scotoductus                          PVRREF.....................................................
UJG41740.1_Candidatus_Heimdallarchaeum_aukensis     FLKNIFFQYIFEKALILYLSELDFSNIDFLPELHNLSADFRI.......LLRKMGYK..
MFW9996705_Candidatus_Odinarchaeota_archaeon        TIQHILDVLFKREASKLRTSGIFFNRTHSVPGFREKSIQIKE.......RLTLLGLN..
MCH8915407.1_Thaumarchaeota_archaeon                PLKNTFALIGSSRE..................YQQFDKNFTVESKGE............
MHA1972560.1_Candidatus_Hodarchaeales_archaeon      GLEEEFLKDVGRY....IPNFILFDI.................................
NHI95076.1_Candidatus_Lokiarchaeota_archaeon        AIRNQFISE..................................................
MEM2144305.1_Candidatus_Jordarchaeaceae_archaeon    EVKYKLVQFQQEFE.............................................
AHB41284.1_candidate_division_SR1_bacterium         PLKNTFGHLHNTRN..................FYHLKKDIAQHNLRD............
HEV2339438.1_Patescibacteria_group_bacterium        KLKQTFALMGSFRT..................YNMFDGNYPVNAKEFT...........
MBL9158386.1_Verrucomicrobiales_bacterium           RLAPAIADSLQRLE.............................................
MGB8583703.1_Candidatus_Sulfotelmatobacter_sp.      APLQAFGPVRLINP.............................................
MBI3627515.1_Candidatus_Sungbacteria_bacterium      AAETKFLETLKQW....IPNFILFNS.................................
_213105201.1_Candidatus_Protochlamydia_amoebophila  SIQQV......................................................
CDRQKP010002909.1_Symbiodinium_sp.                  KLKETVLNS..KN....FPNFIFFEE.................................
JAUKPS010110286.1_Chlorellidium_tetrabotrys         TLAERLRDALLKE....LPKFILFSS.................................
XP_004364654.1_Capsaspora_owczarzaki                KIRFAFVQPFYQW..............................................
KJE91799.1_Capsaspora_owczarzaki                    KVRYAFVQPSFLW..............................................
XP_048575736.1_Nematostella_vectensis               PQKSFLINRREVPY...ASEAITDGEVSTNTSCASLDDL....................
A0A6P8J3J2_Actinia_tenebrosa                        PQHSFLLNVIKVPY...IAEGLL.....RNASCSSLESV....................
XP_022780642.1_Stylophora_pistillata                PDQKFLRKINFDVH.................VECALDAT....................
A0A9X0CFV3_Desmophyllum_pertusum                    PDEAFLHKVKLDNP.................NESTMDAI....................
A0A6S7HLM0_Paramuricea_clavata                      PDESILVST......................GCNFLPDL....................
XP_046856289.1_Xenia_sp.                            PDESKLIST......................GCHFLPDL....................
A0A813MIX4_Adineta_steineri                         ...........................................................
UJR19038.1_Adineta_vaga                             TIEDS......................................................
JAXIUW010014154.1_Cephalothrix_simula               NLTEAFVQVFQGQV.............................................
A0A8B8DK01_Crassostrea_virginica                    DIEERYVATFPLRG.............................................
XP_070564142.1_Ptychodera_flava                     DDSDEVLSSVAGVK.....SFL.............FDWRYAHKIKNNATEILRRASRPH
ACQM01009845.1_Saccoglossus_kowalevskii             EVTNEPLSTLLRVP.....ENIIIPEMKCTTKCDERDVNYPYNRLNMVKECITHGVK..
XP_078582628.1_Branchiostoma_floridae_japonicum     ...........................................................
JALCYT010000124.1_Branchiostoma_belcheri            ...........................................................
KAJ3014353.1_Thoreauomyces_humboldtii               GQTAKLLGPLSPR..............................................
KAJ3188706.1_Gaertneriomyces_sp._JEL0708            SISRK......................................................
XP_002110806.1_Trichoplax_adhaerens                 ...........................................................
RDD37009.1_Trichoplax_sp._H2                        ...........................................................
A0A8W8L770_Magallana_gigas                          TIENKYVSTLPLRG.............................................
A0A8W8L770_Magallana_gigas                          ℓℓℓ  ➔
                                                    α5
```

*E8PLM2_Thermus_scotoductus*

```
E8PLM2_Thermus_scotoductus                          ...........................................................
UJG41740.1_Candidatus_Heimdallarchaeum_aukensis     .IE..........IYNYYKYFGNFISKIIRQKVVILREDR..GLLEN...........G
MFW9996705_Candidatus_Odinarchaeota_archaeon        .TE............NNFSDLPQAIILTLREKTGILYEDR..GLLNSL..........G
MCH8915407.1_Thaumarchaeota_archaeon                ..................RY.QNLKNKLAEESTIKIESGE...................
MHA1972560.1_Candidatus_Hodarchaeales_archaeon      ........................YNDQIPNKVPIAQLGS...................
NHI95076.1_Candidatus_Lokiarchaeota_archaeon        ...........................................................
MEM2144305.1_Candidatus_Jordarchaeaceae_archaeon    ...........................................................
AHB41284.1_candidate_division_SR1_bacterium         .....................DFISKKNYRENNSL........................
HEV2339438.1_Patescibacteria_group_bacterium        ..................RT.AELKTKDVNENTRASSNEE...................
MBL9158386.1_Verrucomicrobiales_bacterium           ...........................................................
MGB8583703.1_Candidatus_Sulfotelmatobacter_sp.      ...........................................................
MBI3627515.1_Candidatus_Sungbacteria_bacterium      ........................FEDIFPNKIPFAELEK...................
_213105201.1_Candidatus_Protochlamydia_amoebophila  ...........................................................
CDRQKP010002909.1_Symbiodinium_sp.                  .........................EDEIKDTYDLDSIGD...................
JAUKPS010110286.1_Chlorellidium_tetrabotrys         ........................FEDVFPNEIPFGELKS...................
XP_004364654.1_Capsaspora_owczarzaki                ..........................GLYSKEVQ.........................
KJE91799.1_Capsaspora_owczarzaki                    ..........................GEHQRQED.........................
XP_048575736.1_Nematostella_vectensis               ........................FHSNMPRNASMQS.....NISMEMTAHDQSD....
A0A6P8J3J2_Actinia_tenebrosa                        ........................FQCNFANKVALEPTTE..GVAMETVEGLQSDVDGG
XP_022780642.1_Stylophora_pistillata                .............................EEPLSLASPLP...................
A0A9X0CFV3_Desmophyllum_pertusum                    .............................EQPLSLESSSD...................
A0A6S7HLM0_Paramuricea_clavata                      ...........................................................
XP_046856289.1_Xenia_sp.                            ...........................................................
A0A813MIX4_Adineta_steineri                         ...........................CLSKSDY........GIFESLFPSV.......
UJR19038.1_Adineta_vaga                             ...........................CQEREEQIFHDECHTCIPKNLFPVV.......
JAXIUW010014154.1_Cephalothrix_simula               ......................KTCNVQWSRSERVTKGGDG..................
A0A8B8DK01_Crassostrea_virginica                    .......................IGMVQWTKSDRIK.......................
XP_070564142.1_Ptychodera_flava                     SMEDGQTIYKKIITYTKKKIRKSFSTGSFTPPMQMTSADKG..................
ACQM01009845.1_Saccoglossus_kowalevskii             .VEGQKEFIPMVVKFLQDKV....LSRTSTDGSGMTAPNVG..................
XP_078582628.1_Branchiostoma_floridae_japonicum     .........................DGQETEESDGNNEDI...................
JALCYT010000124.1_Branchiostoma_belcheri            .........................DGQETDSEGCEDIDC...................
KAJ3014353.1_Thoreauomyces_humboldtii               ..........................GSSTGDEDLDEDAP...................
KAJ3188706.1_Gaertneriomyces_sp._JEL0708            .............................SKRSALAVPDPP..................
XP_002110806.1_Trichoplax_adhaerens                 .......................YYSSNFRKNEDL........................
RDD37009.1_Trichoplax_sp._H2                        .......................YYSSNFRKNEDL........................
A0A8W8L770_Magallana_gigas                          .......................IGMVQWTRSEKIKKEYK...................
A0A8W8L770_Magallana_gigas                                                 ℓℓℓ   TT TT  ℓℓℓℓ...................
```

*E8PLM2_Thermus_scotoductus*

```
                                                                                                                   ℓℓℓℓ
                                                                                                                170
                                                                                                                   .
E8PLM2_Thermus_scotoductus                          .....................................................KLQDNF
UJG41740.1_Candidatus_Heimdallarchaeum_aukensis     F..................SY......................PWRENITNVLTLERIK
MFW9996705_Candidatus_Odinarchaeota_archaeon        LK...............................................ISTFGLKETI
MCH8915407.1_Thaumarchaeota_archaeon                ................................................PSALTYVKLKL
MHA1972560.1_Candidatus_Hodarchaeales_archaeon      ...........................................................
NHI95076.1_Candidatus_Lokiarchaeota_archaeon        ........................................................GRS
MEM2144305.1_Candidatus_Jordarchaeaceae_archaeon    ........................................................HTL
AHB41284.1_candidate_division_SR1_bacterium         .................................................LGYALCIKKL
HEV2339438.1_Patescibacteria_group_bacterium        ................................................PVAFMYVKKEL
MBL9158386.1_Verrucomicrobiales_bacterium           ..........................................................H
MGB8583703.1_Candidatus_Sulfotelmatobacter_sp.      ........................................HRVPQPRQPLQSVSNLTGD
MBI3627515.1_Candidatus_Sungbacteria_bacterium      ...........................................................
_213105201.1_Candidatus_Protochlamydia_amoebophila  ......................................................AKLTF
CDRQKP010002909.1_Symbiodinium_sp.                  ...........................................................
JAUKPS010110286.1_Chlorellidium_tetrabotrys         ...........................................................
XP_004364654.1_Capsaspora_owczarzaki                .................................................GSSKHLLTTG
KJE91799.1_Capsaspora_owczarzaki                    .................................................TGDKHLLTTG
XP_048575736.1_Nematostella_vectensis               .....EGVRLRHASIDSCRSSLASVLL.......KRPAWRRQSSYGGLAWS..NYHELK
A0A6P8J3J2_Actinia_tenebrosa                        IIGMMQGDGGKIFKTGHCETENASPKLERRQTAGQRHSFKRMSSFGGLAWS..NYHELK
XP_022780642.1_Stylophora_pistillata                ..................................KDMIHNKPGTFTNNSWETLSYQDIK
A0A9X0CFV3_Desmophyllum_pertusum                    ..................................TDFMQTKHAHFVNSNWDTLRYRDVK
A0A6S7HLM0_Paramuricea_clavata                      .......................................KTPVSNEDNSNECLNPKNPQ
XP_046856289.1_Xenia_sp.                            .......................................K...ESNNNPIEYLNLKHPQ
A0A813MIX4_Adineta_steineri                         ......................................................KAYDT
UJR19038.1_Adineta_vaga                             ......................................................VPFDS
JAXIUW010014154.1_Cephalothrix_simula               .................................................TNNYRETDQN
A0A8B8DK01_Crassostrea_virginica                    .................................................EDHYAEACKR
XP_070564142.1_Ptychodera_flava                     ........................................SRQDYHRRVSHLCATGKID
ACQM01009845.1_Saccoglossus_kowalevskii             ........................................TRTDCLRYIQQKIKRGAIK
XP_078582628.1_Branchiostoma_floridae_japonicum     ..............................................KISQNVYKICVDN
JALCYT010000124.1_Branchiostoma_belcheri            .................................................KGVFKIRVDN
KAJ3014353.1_Thoreauomyces_humboldtii               ...........................................SLSEKLSERLWDGRHL
KAJ3188706.1_Gaertneriomyces_sp._JEL0708            ........................................FDLLIPGSLVNAMRTGNGQ
XP_002110806.1_Trichoplax_adhaerens                 ...........................................................
RDD37009.1_Trichoplax_sp._H2                        ..................................................GQFRYFNKD
A0A8W8L770_Magallana_gigas                          .................................................ESNYKMACDR
A0A8W8L770_Magallana_gigas                          .................................................ℓℓℓℓℓℓℓℓℓ
```

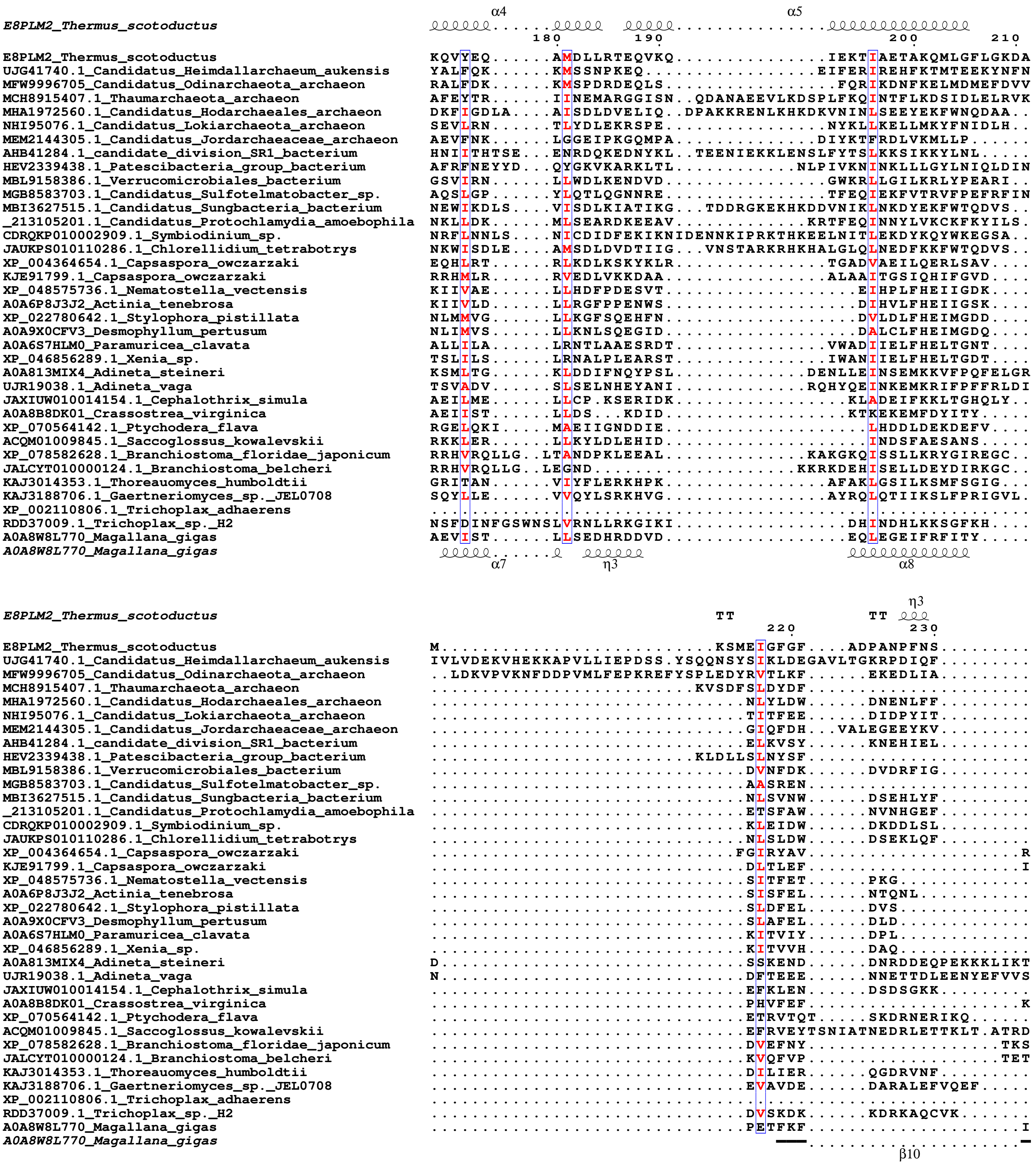
E8PLM2_Thermus_scotoductus
α4
α5
180
190
200
210
E8PLM2_Thermus_scotoductus KQVYEQ......AMDLLRTEQVKQ...............IEKTIAETAKQMLGFLGKDA
UJG41740.1_Candidatus_Heimdallarchaeum_aukensis YALFQK......KMSSNPKEQ.................EIFERIREHFKTMTEEKYNFN
MFW9996705_Candidatus_Odinarchaeota_archaeon RALFDK......KMSPDRDEQLS.................FQRIKDNFKELMDMEFDVV
MCH8915407.1_Thaumarchaeota_archaeon AFEYTR......IINEMARGGISN..QDANAEEVLKDSPLFKQINTFLKDSIDLELRVK
MHA1972560.1_Candidatus_Hodarchaeales_archaeon DKFIGDLA...AISDLDVELIQ..DPAKKRENLKHKDKVNINLSEEYEKFWNQDAA..
NHI95076.1_Candidatus_Lokiarchaeota_archaeon SEVLRN......TLYDLEKRSPE................NYKLLKELLMKYFNIDLH..
MEM2144305.1_Candidatus_Jordarchaeaceae_archaeon AEVFNK......LGGEIPKGQMPA..............DIYKTFRDLVKMLLP......
AHB41284.1_candidate_division_SR1_bacterium HNIITHTSE...ENRDQKEDNYKL..TEENIEKKLENSLFYTSLKKSIKKYLNL.....
HEV2339438.1_Patescibacteria_group_bacterium AFRFNEYYD...QYGKVKARKLTL............NLPIVKNINKLLLGYLNIQLDIN
MBL9158386.1_Verrucomicrobiales_bacterium GSVIRN......LLWDLKENDVD................GWKRLLGILKRLYPEARI..
MGB8583703.1_Candidatus_Sulfotelmatobacter_sp. AQSLGP......YLQTLQGNNRE................TFEQIEKFVTRVFPEFRFIN
MBI3627515.1_Candidatus_Sungbacteria_bacterium NEWIKDLS....VISDLKIATIKG...TDDRGKEKHKDDVNIKLNKDYEKFWTQDVS..
_213105201.1_Candidatus_Protochlamydia_amoebophila NKLLDK......MLSEARDKEEAV.............KRTFEQINNYLVKCKFKYILS.
CDRQKP010002909.1_Symbiodinium_sp. NRFLNNLS....NICDIDFEKIKNIDENNKIPRKTHKEELNITLEKDYKQYWKEGSA..
JAUKPS010110286.1_Chlorellidium_tetrabotrys NKWISDLE....AMSDLDVDTIIG...VNSTARKRHKHALGLQLNEDFKKFWTQDVS..
XP_004364654.1_Capsaspora_owczarzaki EQHLRT......RLKDLKSKYKLR...............TGADVAEILQERLSAV....
KJE91799.1_Capsaspora_owczarzaki RRHMLR......RVEDLVKKDAA................ALAAITGSIQHIFGVD....
XP_048575736.1_Nematostella_vectensis KIIVAE......LLHDFPDESVT...................EIHPLFHEIIGDK....
A0A6P8J3J2_Actinia_tenebrosa KIIVLD......LLRGFPPENWS...................DIHVLFHEIIGSK....
XP_022780642.1_Stylophora_pistillata NLMMVG......LLKGFSQEHFN...................DVLDLFHEIMGDD....
A0A9X0CFV3_Desmophyllum_pertusum NLIMVS......LLKNLSQEGID...................DALCLFHEIMGDQ....
A0A6S7HLM0_Paramuricea_clavata ALLILA......LRNTLAAESRDT...............VWADIIELFHELTGNT....
XP_046856289.1_Xenia_sp. TSLILS......LRNALPLEARST...............IWANIIELFHELTGDT....
A0A813MIX4_Adineta_steineri KSMLTG......KLDDIFNQYPSL.............DENLLEINSEMKKVFPQFELGR
UJR19038.1_Adineta_vaga TSVADV......SLSELNHEYANI.............RQHYQEINKEMKRIFPFFRLDI
JAXIUW010014154.1_Cephalothrix_simula AEILME......LLCP.KSERIDK.................KLADEIFKKLTGHQLY..
A0A8B8DK01_Crassostrea_virginica AEIIST......LLDS...KDID..................KTKEKEMFDYITY.....
XP_070564142.1_Ptychodera_flava RGELQKI.....MAEIIGNDDIE....................LHDDLDEKDEFV....
ACQM01009845.1_Saccoglossus_kowalevskii RKKLER......LLKYLDLEHID....................INDSFAESANS.....
XP_078582628.1_Branchiostoma_floridae_japonicum RRHVRQLLG..LTANDPKLEEAL..............KAKGKQISSLLKRYGIREGC..
JALCYT010000124.1_Branchiostoma_belcheri RRHVRQLLG..LEGND....................KKRKDEHISELLDEYDIRKGC..
KAJ3014353.1_Thoreauomyces_humboldtii GRITAN......VIYFLERKHPK................AFAKLGSILKSMFSGIG...
KAJ3188706.1_Gaertneriomyces_sp._JEL0708 SQYLLE......VVQYLSRKHVG................AYRQLQTIIKSLFPRIGVL.
XP_002110806.1_Trichoplax_adhaerens ...........................................................
RDD37009.1_Trichoplax_sp._H2 NSFDINFGSWNSLVRNLLRKGIKI.................DHINDHLKKSGFKH....
A0A8W8L770_Magallana_gigas AEVIST......LLSEDHRDDVD..................EQLEGEIFRFITY.....
A0A8W8L770_Magallana_gigas
α7
η3
α8
E8PLM2_Thermus_scotoductus
TT
TT
η3
220
230
E8PLM2_Thermus_scotoductus M...........................KSMEIGFGF...ADPANPFNS.........
UJG41740.1_Candidatus_Heimdallarchaeum_aukensis IVLVDEKVHEKKAPVLLIEPDSS.YSQQNSYSIKLDEGAVLTGKRPDIQF.........
MFW9996705_Candidatus_Odinarchaeota_archaeon ..LDKVPVKNFDDPVMLFEPKREFYSPLEDYRVTLKF......EKEDLIA.........
MCH8915407.1_Thaumarchaeota_archaeon ..........................KVSDFSLDYDF......................
MHA1972560.1_Candidatus_Hodarchaeales_archaeon ...............................NLYLDW......DNENLFF.........
NHI95076.1_Candidatus_Lokiarchaeota_archaeon ...............................TITFEE......DIDPYIT.........
MEM2144305.1_Candidatus_Jordarchaeaceae_archaeon ...............................GIQFDH...VALEGEEYKV.........
AHB41284.1_candidate_division_SR1_bacterium ...............................ELKVSY......KNEHIEL.........
HEV2339438.1_Patescibacteria_group_bacterium ..........................KLDLLSLNYSF......................
MBL9158386.1_Verrucomicrobiales_bacterium ...............................DVNFDK......DVDRFIG.........
MGB8583703.1_Candidatus_Sulfotelmatobacter_sp. ...............................AASREN......................
MBI3627515.1_Candidatus_Sungbacteria_bacterium ...............................NLSVNW......DSEHLYF.........
_213105201.1_Candidatus_Protochlamydia_amoebophila ...............................ETSFAW......NVNHGEF.........
CDRQKP010002909.1_Symbiodinium_sp. ...............................KLEIDW......DKDDLSL.........
JAUKPS010110286.1_Chlorellidium_tetrabotrys ...............................NLSLDW......DSEKLQF.........
XP_004364654.1_Capsaspora_owczarzaki ..............................FGIRYAV.....................R
KJE91799.1_Capsaspora_owczarzaki ...............................DLTLEF.....................I
XP_048575736.1_Nematostella_vectensis ...............................SITFET......PKG.............
A0A6P8J3J2_Actinia_tenebrosa ...............................SISFEL......NTQNL...........
XP_022780642.1_Stylophora_pistillata ...............................SLDFEL......DVS.............
A0A9X0CFV3_Desmophyllum_pertusum ...............................SLAFEL......DLD.............
A0A6S7HLM0_Paramuricea_clavata ...............................KITVIY......DPL.............
XP_046856289.1_Xenia_sp. ...............................KITVVH......DAQ.............
A0A813MIX4_Adineta_steineri D..............................SSKEND......DNRDDEQPEKKKLIKT
UJR19038.1_Adineta_vaga N..............................DFTEEE......NNETTDLEENYEFVVS
JAXIUW010014154.1_Cephalothrix_simula ...............................EFKLEN......DSDSGKK.........
A0A8B8DK01_Crassostrea_virginica ...............................PHVFEF.....................K
XP_070564142.1_Ptychodera_flava ...............................ETRVTQT.....SKDRNERIKQ......
ACQM01009845.1_Saccoglossus_kowalevskii ...............................EFRVEYTSNIATNEDRLETTKLT.ATRD
XP_078582628.1_Branchiostoma_floridae_japonicum ...............................DVEFNY...................TKS
JALCYT010000124.1_Branchiostoma_belcheri ...............................KVQFVP...................TET
KAJ3014353.1_Thoreauomyces_humboldtii ...............................DILIER......QGDRVNF.........
KAJ3188706.1_Gaertneriomyces_sp._JEL0708 ...............................EVAVDE......DARALEFVQEF.....
XP_002110806.1_Trichoplax_adhaerens ...........................................................
RDD37009.1_Trichoplax_sp._H2 ...............................DVSKDK......KDRKAQCVK.......
A0A8W8L770_Magallana_gigas ...............................PETFKF.....................I
A0A8W8L770_Magallana_gigas
β10

```
                                                      β9                                                α6
E8PLM2_Thermus_scotoductus
                                                                                   240          250          260
E8PLM2_Thermus_scotoductus                        .....LRLQYR........ESD...............LTLPGDELGLGIQSAIVVGIFE
UJG41740.1_Candidatus_Heimdallarchaeum_aukensis   ..............HDL..QNN...............KAFRLNCSPGGAYEILMVLCSI
MFW9996705_Candidatus_Odinarchaeota_archaeon      HEP..GLIFYS........TKD.............ERMSNTIDQAPGGAYEIIMVLTVV
MCH8915407.1_Thaumarchaeota_archaeon              .......SFVD........KNG...............IEIQIGELSSGQRGIIHFIFSI
MHA1972560.1_Candidatus_Hodarchaeales_archaeon    ........WIK........ENE...............EYYEPKQRSRGRQWHLSFYVRV
NHI95076.1_Candidatus_Lokiarchaeota_archaeon      .......CEYK........EEN...............TSLDLICGGSGFLQILHLLTFI
MEM2144305.1_Candidatus_Jordarchaeaceae_archaeon  .......YFIN........RMG...............IRVDFDQLSSGEKDIIAMLFPF
AHB41284.1_candidate_division_SR1_bacterium       .......YFIN........TTG...............QEQTLHSISNGEQSLLIILLTI
HEV2339438.1_Patescibacteria_group_bacterium      .......KFID........EKK...............NEVQYSDLSAGQKAIIHFIFCL
MBL9158386.1_Verrucomicrobiales_bacterium         .......SGYS........DKV.............LERNLDVVVSGTGFQQVLQIFTGV
MGB8583703.1_Candidatus_Sulfotelmatobacter_sp.    .NQ..VSIDLT........ERS.............TGRKIPLENCGTGVEQTIILATFV
MBI3627515.1_Candidatus_Sungbacteria_bacterium    ........WVK........EDR...............YPYEPSLRSKGRQWHLAFYIKV
_213105201.1_Candidatus_Protochlamydia_amoebophila .KD..VNLTFA........NKD..........LQMKYSTIPLSNISSGERIVLLILLWR
CDRQKP010002909.1_Symbiodinium_sp.                ........YIK........EGK...............NTFLPKQRSRGKNWFLSFYVLL
JAUKPS010110286.1_Chlorellidium_tetrabotrys       ........WIQ........EGD...............EFYPPEVRSQGRRWHLAFYIRV
XP_004364654.1_Capsaspora_owczarzaki              IAS..IVVYVQ........EEL..P.......DGTMGAELEIGACSSSLQKVMALYVLF
KJE91799.1_Capsaspora_owczarzaki                  DDP..NRLFVK........ESG...............VQLDIGMCAASLQKVVAIYVLF
XP_048575736.1_Nematostella_vectensis             ..E..DVFQVM........DNK...............TKRVMRRIPEGLFSAFIIAALL
A0A6P8J3J2_Actinia_tenebrosa                      .PQ..ESFKIV........DTK...............TGRVMRRIPEGLFTAFIIAALL
XP_022780642.1_Stylophora_pistillata              .SN..IPV.IL........NKR...............TQRIIQQVSEGTFSAFVVAALV
A0A9X0CFV3_Desmophyllum_pertusum                  .GD..IPIRIL........DKG...............TRRIIQRISEGTFSTFVVAALV
A0A6S7HLM0_Paramuricea_clavata                    ..S..GAVRMK........QQY.............GHCETIRDDLPEGFFHAFLIAFLV
XP_046856289.1_Xenia_sp.                          ..S..NLIHVR........QQY.............GDCHIIRDDLPEGFFHAFLIAFLI
A0A813MIX4_Adineta_steineri                       TSK..STFFIRLT......SRNY..............LFSDISDCSDGFLRYLTLLIAI
UJR19038.1_Adineta_vaga                           IRQYLSPGYYSVT........................EFSDVTNCGEGILRCLYIIIKI
JAXIUW010014154.1_Cephalothrix_simula             .......IKVK........YNK...............RYHPLLKTPEGIIAAKEIALIV
A0A8B8DK01_Crassostrea_virginica                  KKG..EEIMVS........RAD..........DKIEYRYFPLLKTSEGVLEAKATSLLL
XP_070564142.1_Ptychodera_flava                   IRK..IQTSYN........DNSTEPATTSYCWDKITKQKFYIDQLPSGLASAFEIASQL
ACQM01009845.1_Saccoglossus_kowalevskii           TKT..NTVSVK.......NEVD.....SAKCEDGNTGKTFNIKQLPSGDASTYEAAAML
XP_078582628.1_Branchiostoma_floridae_japonicum   SKT..LKIAVTVSEQGVGRNQD...............VSLPLSQCDAGLQQVIIIMLMV
JALCYT010000124.1_Branchiostoma_belcheri          QNE..LQIVVV........EQG...............VTLPLFQYDAGLQQVIIIMLMV
KAJ3014353.1_Thoreauomyces_humboldtii             ........QFS........RTE.............TTTTSSWVREGSGVAYVSSVFAAV
KAJ3188706.1_Gaertneriomyces_sp._JEL0708          ...................ESG..........RKRKQIRRPWHEEGSGVCYAVTLFAAI
XP_002110806.1_Trichoplax_adhaerens               ...................EKT.....................................
RDD37009.1_Trichoplax_sp._H2                      .....RDFYYD........KSDV....................QISPGESLELLSALWL
A0A8W8L770_Magallana_gigas                        KKD..GLLYVQ........HNE...............SEFPLLKTSEGILEAKLTSLLL
A0A8W8L770_Magallana_gigas                        →T..T———........▶TT                  ——→ η4       α9
                                                          β11                          β12

                                                                                                 β10
E8PLM2_Thermus_scotoductus
                                                                                        270          280
E8PLM2_Thermus_scotoductus                        AFRQL....................................GEKIGTVIIEEPEMYLHP
UJG41740.1_Candidatus_Heimdallarchaeum_aukensis   YGFEA........................................DTIILDEPGKTLHP
MFW9996705_Candidatus_Odinarchaeota_archaeon      ELSKT........................................TTILLDEPGKTLHP
MCH8915407.1_Thaumarchaeota_archaeon              FGYDL......................................ENGLMIIDEPELHIHP
MHA1972560.1_Candidatus_Hodarchaeales_archaeon    TARSR....................................EDVSNIILIDEPGLFLHA
NHI95076.1_Candidatus_Lokiarchaeota_archaeon      LTKST........................................SIILLDEPDAHLHP
MEM2144305.1_Candidatus_Jordarchaeaceae_archaeon  IEKEV......................ENE..LSQTKSEKIPHEDLVIMIDSPEAYLHP
AHB41284.1_candidate_division_SR1_bacterium       YGFDL......................................KDGILIIDEPELHFHP
HEV2339438.1_Patescibacteria_group_bacterium      YGYDI......................................EKGLLIIDEPETHLHP
MBL9158386.1_Verrucomicrobiales_bacterium         LSQGS........................................NIVLLDEPDAHLHG
MGB8583703.1_Candidatus_Sulfotelmatobacter_sp.    LT.......................................TPKPGTILLDEPHSYLHP
MBI3627515.1_Candidatus_Sungbacteria_bacterium    SARAS....................................ENTPNIILIDEPGLFLHA
_213105201.1_Candidatus_Protochlamydia_amoebophila FDQRN....................................IQKESVILLDEPDAHLHP
CDRQKP010002909.1_Symbiodinium_sp.                CSRFD....................................ENKKNIILIDEPGQYLHP
JAUKPS010110286.1_Chlorellidium_tetrabotrys       AARSR....................................DGVSSVILIDEPGLYLHA
XP_004364654.1_Capsaspora_owczarzaki              FLLAFSTAADAAPSGEIPSPASSPDVSEESKGLPDPAAWKSPSTQRILLVDELEALLYE
KJE91799.1_Capsaspora_owczarzaki                  HLLRVSTAGEA................EGRQ.....DCDDNEATQRILLIDEPEALLYD
XP_048575736.1_Nematostella_vectensis             VKPST........................................RNVILDEPTRGMHP
A0A6P8J3J2_Actinia_tenebrosa                      VKPGS........................................RNVILDEPTRGMHP
XP_022780642.1_Stylophora_pistillata              VCPFT........................................HTLIFDEVARGMHP
A0A9X0CFV3_Desmophyllum_pertusum                  VQPFS........................................RTVIFDEVCRGMHP
A0A6S7HLM0_Paramuricea_clavata                    VNPTV........................................KTILLDEPTRGMHP
XP_046856289.1_Xenia_sp.                          LNPAN........................................STVLLDEPTRGMHP
A0A813MIX4_Adineta_steineri                       LCQPL.......................................HSIITFDEPDAHIFP
UJR19038.1_Adineta_vaga                           CTVKE.......................................NSIITIDEPDAHVFP
JAXIUW010014154.1_Cephalothrix_simula             AHKEF........................................KTILFEEIDQGMNP
A0A8B8DK01_Crassostrea_virginica                  ALNNI........................................QTLCLEEPDKSMHP
XP_070564142.1_Ptychodera_flava                   ARKEV........................................KTLLLDEPSQCMHI
ACQM01009845.1_Saccoglossus_kowalevskii           AQINV........................................QTIIFDDPDMCMHP
XP_078582628.1_Branchiostoma_floridae_japonicum   LHVQE..................................KEGKKPRHLLLDEPVSLFHG
JALCYT010000124.1_Branchiostoma_belcheri          LYVQG..................................KEGKRHVTLLLDEPVSLFHG
KAJ3014353.1_Thoreauomyces_humboldtii             LFDQV......V......RFSS.......IQRL...PQAGDPEPVHILALDEPSSPLHP
KAJ3188706.1_Gaertneriomyces_sp._JEL0708          IFDSV......V......RFQSRADGAENSTGM...DAAPLQQALHILAVDEPSTPFHP
XP_002110806.1_Trichoplax_adhaerens               ......................................DKTEKKKAVYLLDEPDAHLHT
RDD37009.1_Trichoplax_sp._H2                      FEIET.................................DTKSNKRAVYLLDEPDAHLHT
A0A8W8L770_Magallana_gigas                        AHKHI........................................QTLCLEDPDRGMHP
A0A8W8L770_Magallana_gigas
                                                                                                 β13
```

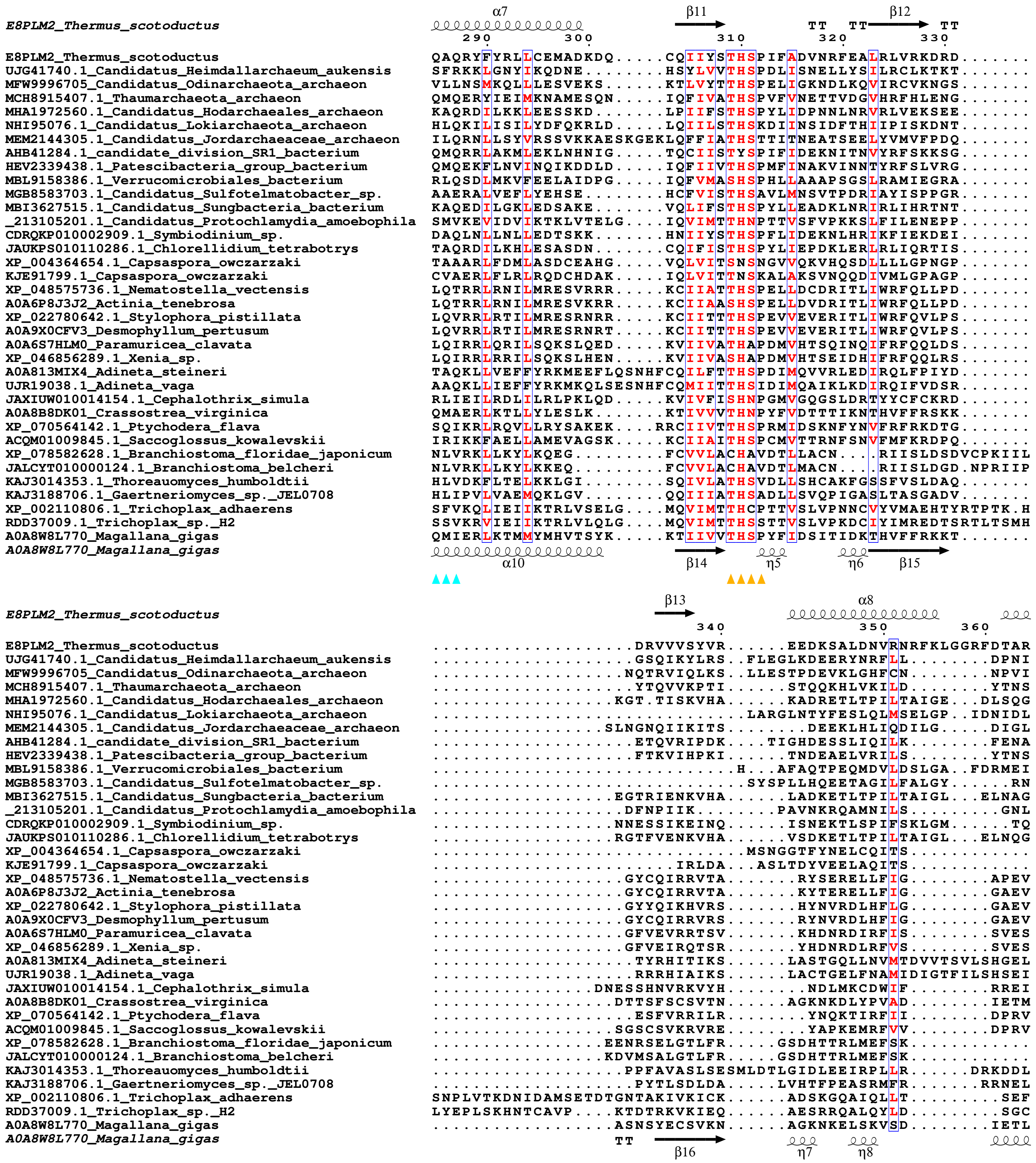

E8PLM2_Thermus_scotoductus
α7
β11
TT
TT
β12
TT
290
300
310
320
330
E8PLM2_Thermus_scotoductus QAQRYFYRLLCEMADKDQ.....CQIIYSTHSPIFADVNRFEALRLVRKDRD......
UJG41740.1_Candidatus_Heimdallarchaeum_aukensis SFRKKLGNYIKQDNE........HSYLVVTHSPDLISNELLYSILRCLKTKT......
MFW9996705_Candidatus_Odinarchaeota_archaeon VLLNSMKQLLLESVEKS......KTLVYTTHSPELIGKNDLKQVIRCVKNGS......
MCH8915407.1_Thaumarchaeota_archaeon QMQERYIEIMKNAMESQN.....IQFIVATHSPVFVNETTVDGVHRFHLENG......
MHA1972560.1_Candidatus_Hodarchaeales_archaeon KAQRDILKKLEESSKD.......LPIIFSTHSPYLIDPNNLNRVRLVEKSEE......
NHI95076.1_Candidatus_Lokiarchaeota_archaeon HLQKILISILYDFQKRLD.....LQIILSTHSKDIINSIDFTHIIPISKDNT......
MEM2144305.1_Candidatus_Jordarchaeaceae_archaeon ILQRNLLSYVRSSVKKAESKGEKLQFFIATHSTTITNEATSEELYVMVFPDQ......
AHB41284.1_candidate_division_SR1_bacterium QMQRRLAKMLEKLNHNIG.....TQCIISTYSPIFIDEKNITNVYRFSKKSG......
HEV2339438.1_Patescibacteria_group_bacterium QMQEKFLNVINQIKDDLD.....IQFIIVTHSPMFINAKVINNTYRFSLVRG......
MBL9158386.1_Verrucomicrobiales_bacterium RLQSDLMKVFEELAIDPG.....IQFVMASHSPHLLAAAPSGSLRAMIEGRA......
MGB8583703.1_Candidatus_Sulfotelmatobacter_sp. AAERALVEFLYEHSE........HCFVISTHSAVLMNSVTPDRIAYISPPGR......
MBI3627515.1_Candidatus_Sungbacteria_bacterium KAQEDILGKLEDSAKE.......VQLIFSTHSPYLLEADKLNRIRLIHRTNT......
_213105201.1_Candidatus_Protochlamydia_amoebophila SMVKEVIDVIKTKLVTELG....IQVIMTTHNPTTVSFVPKKSLFILENEPP......
CDRQKP010002909.1_Symbiodinium_sp. DAQLNLLNLLEDTSKK.......HNIIYSTHSPFLIDEKNLHRIKFIEKDEI......
JAUKPS010110286.1_Chlorellidium_tetrabotrys TAQRDILKHLESASDN.......CQIFISTHSPYLIEPDKLERLRLIQRTIS......
XP_004364654.1_Capsaspora_owczarzaki TAAARLFDMLASDCEAHG.....VQLVITSNSNGVVQKVHQSDLLLLGPNGP......
KJE91799.1_Capsaspora_owczarzaki CVAERLFLRLRQDCHDAK.....IQLVITTNSKALAKSVNQQDIVMLGPAGP......
XP_048575736.1_Nematostella_vectensis LQTRRLRNILMRESVRRR.....KCIIATTHSPELLDCDRITLIWRFQLLPD......
A0A6P8J3J2_Actinia_tenebrosa LQTRRLRNILMRESVKRR.....KCIIAASHSPELLDVDRITLIWRFQLLPD......
XP_022780642.1_Stylophora_pistillata LQVRRLRTILMRESRNRR.....KCIITTTHSPEVVEVERITLIWRFQVLPS......
A0A9X0CFV3_Desmophyllum_pertusum LQVRRLRTILMRESRNRT.....KCIITTTHSPEVVEVERITLIWRFQVLPS......
A0A6S7HLM0_Paramuricea_clavata LQIRRLQRILSQKSLQED.....KVIIVATHAPDMVHTSQINQIFRFQQLDS......
XP_046856289.1_Xenia_sp. LQIRRLRRILSQKSLHEN.....KVIIVASHAPDMVHTSEIDHIFRFQQLRS......
A0A813MIX4_Adineta_steineri TAQKLLVEFFYRKMEEFLQSNHFCQILFTTHSPDIMQVVRLEDIRQLFPIYD......
UJR19038.1_Adineta_vaga AAQKLLIEFFYRKMKQLSESNHFCQMIITTHSIDIMQAIKLKDIRQIFVDSR......
JAXIUW010014154.1_Cephalothrix_simula RLIEILRDLILRLPKLQD.....KVIVFISHNPGMVGQGSLDRTYYCFCKRD......
A0A8B8DK01_Crassostrea_virginica QMAERLKTLLYLESLK.......KTIVVVTHNPYFVDTTTIKNTHVFFRSKK......
XP_070564142.1_Ptychodera_flava SQIKRLRQVLLRYSAKEK....RRCIIVTTHSPRMIDSKNFYNVFRFRKDTG......
ACQM01009845.1_Saccoglossus_kowalevskii IRIKKFAELLAMEVAGSK.....KCIIAITHSPCMVTTRNFSNVFMFKRDPQ......
XP_078582628.1_Branchiostoma_floridae_japonicum NLVRKLLKYLKQEG.........FCVVLACHAVDTLLACN....RIISLDSDVCPKIIL
JALCYT010000124.1_Branchiostoma_belcheri NLVRKLLKYLKKEQ.........FCVVLACHAVDTLMACN....RIISLDGD.NPRIIP
KAJ3014353.1_Thoreauomyces_humboldtii HLVDKFLTELKKLGI........SQIVLATHSVDLLSHCAKFGSSFVSLDAQ......
KAJ3188706.1_Gaertneriomyces_sp._JEL0708 HLIPVLVAEMQKLGV........QQIIIATHSADLLSVQPIGASLTASGADV......
XP_002110806.1_Trichoplax_adhaerens SFVKQLIEIIKTRLVSELG....MQVIMTTHCPTTVSLVPNNCVYVMAEHTYRTPTK.H
RDD37009.1_Trichoplax_sp._H2 SSVKRVIEIIKTRLVLQLG....MQVIMTTHSSTTVSLVPKDCIYIMREDTSRTLTSMH
A0A8W8L770_Magallana_gigas QMIERLKTMMYMHVTSYK.....KTIIVVTHSPYFIDSITIDKTHVFFRKKT......
A0A8W8L770_Magallana_gigas
α10
β14
η5
η6
β15
E8PLM2_Thermus_scotoductus
β13
α8
340
350
360
E8PLM2_Thermus_scotoductus ....................DRVVVSYVR......EEDKSALDNVRNRFKLGGRFDTAR
UJG41740.1_Candidatus_Heimdallarchaeum_aukensis ....................GSQIKYLRS..FLEGLKDEERYNRFLL........DPNI
MFW9996705_Candidatus_Odinarchaeota_archaeon ....................NQTRVIQLKS..LLESTPDEVKLGHFCN........NPVI
MCH8915407.1_Thaumarchaeota_archaeon ....................YTQVVKPTI......STQQKHLVKILD........YTNS
MHA1972560.1_Candidatus_Hodarchaeales_archaeon ..................KGT.TISKVHA......KADRETLTPILTAIGE...DLSQG
NHI95076.1_Candidatus_Lokiarchaeota_archaeon ..............................LARGLNTYFESLQLMSELGP..IDNIDL
MEM2144305.1_Candidatus_Jordarchaeaceae_archaeon ................SLNGNQIIKITS........DEEKLHLIQDILG.....DIGL
AHB41284.1_candidate_division_SR1_bacterium ....................ETQVRIPDK....TIGHDESSLIQILK........FENA
HEV2339438.1_Patescibacteria_group_bacterium ....................FTKVIHPKI......TNDEAELVRILS........YTNS
MBL9158386.1_Verrucomicrobiales_bacterium .............................H...AFAQTPEQMDVLDSLGA...FDRMEI
MGB8583703.1_Candidatus_Sulfotelmatobacter_sp. ..............................SYSPLLHQEETAGILFALGY......RN
MBI3627515.1_Candidatus_Sungbacteria_bacterium ..................EGTRIENKVHA......LADKETLTPILTAIGL...ELNAG
_213105201.1_Candidatus_Protochlamydia_amoebophila ....................DFNPIIK........PAVNKRQAMNILS.........GNL
CDRQKP010002909.1_Symbiodinium_sp. ..................NNESSIKEINQ......ISNEKTLSPIFSKLGM......TQ
JAUKPS010110286.1_Chlorellidium_tetrabotrys ..................RGTFVENKVHA......VSDKETLTPILTAIGL...ELNQG
XP_004364654.1_Capsaspora_owczarzaki ..............................MSNGGTFYNELCQITS............
KJE91799.1_Capsaspora_owczarzaki ........................IRLDA....ASLTDYVEELAQITS............
XP_048575736.1_Nematostella_vectensis ....................GYCQIRRVTA.......RYSERELLFIG........APEV
A0A6P8J3J2_Actinia_tenebrosa ....................GYCQIRRVTA.......KYTERELLFIG........GAEV
XP_022780642.1_Stylophora_pistillata ....................GYYQIKHVRS.......HYNVRDLHFLG........GAEV
A0A9X0CFV3_Desmophyllum_pertusum ....................GYCQIRRVRS.......RYNVRDLHFIG........GAEV
A0A6S7HLM0_Paramuricea_clavata ....................GFVEVRRTSV.......KHDNRDIRFIS........SVES
XP_046856289.1_Xenia_sp. ....................GFVEIRQTSR.......YHDNRDLRFVS........SVES
A0A813MIX4_Adineta_steineri ......................TYRHITIKS......LASTGQLLNVMTDVVTSVLSHGEL
UJR19038.1_Adineta_vaga ......................RRRHIAIKS......LACTGELFNAMIDIGTFILSHSEI
JAXIUW010014154.1_Cephalothrix_simula ................DNESSHNVRKVYH........NDLMKCDWIF........RREI
A0A8B8DK01_Crassostrea_virginica ....................DTTSFSCSVTN......AGKNKDLYPVAD........IETM
XP_070564142.1_Ptychodera_flava ......................ESFVRRILR........YNQKTIRFII........DPRV
ACQM01009845.1_Saccoglossus_kowalevskii ..................SGSCSVKRVRE.........YAPKEMRFVV........DPRV
XP_078582628.1_Branchiostoma_floridae_japonicum .................EENRSELGTLFR.....GSDHTTRLMEFSK............
JALCYT010000124.1_Branchiostoma_belcheri .................KDVMSALGTLFR.....GSDHTTRLMEFSK............
KAJ3014353.1_Thoreauomyces_humboldtii ......................PPFAVASLSESMLDTLGIDLEEIRPLLR......DRKDDL
KAJ3188706.1_Gaertneriomyces_sp._JEL0708 ......................PYTLSDLDA.....LVHTFPEASRMFR........RRNEL
XP_002110806.1_Trichoplax_adhaerens SNPLVTKDNIDAMSETDTGNTAKIVKICK......ADSKGQAIQLLT.........SEF
RDD37009.1_Trichoplax_sp._H2 LYEPLSKHNTCAVP....KTDTRKVKIEQ......AESRRQALQYLD.........SGC
A0A8W8L770_Magallana_gigas ..................ASNSYECSVKN......AGKNKELSKVSD........IETL
A0A8W8L770_Magallana_gigas
TT
β16
η7
η8

```
E8PLM2_Thermus_scotoductus                              η4       β14        α9
                                                                  370       380       390
E8PLM2_Thermus_scotoductus                              NEVLFAKRALLVEGYGDRVAALQLFNQLEVDPDAE........................
UJG41740.1_Candidatus_Heimdallarchaeum_aukensis         KSIFFASNVIFVEGTYDNKIVTTLFSVLENHHIQK........................
MFW9996705_Candidatus_Odinarchaeota_archaeon            KSMFFSKQVCFLEGVTDERFINALVHVLMDTGHKF........................
MCH8915407.1_Thaumarchaeota_archaeon                    SKIFFSNKVILVEGITDEYFFKHFLGDYKVVQELQDKRK....................
MHA1972560.1_Candidatus_Hodarchaeales_archaeon          IRV.DIKNSFVVEGISDYYYLHAFKKILSFDK...........................
NHI95076.1_Candidatus_Lokiarchaeota_archaeon            ALVIKTKKLLFIEGKSDKIIKQLAEISGYNLFVNN........................
MEM2144305.1_Candidatus_Jordarchaeaceae_archaeon        ASLTTGKPILLLEGKSDVEISKLLKPDIEE.............................
AHB41284.1_candidate_division_SR1_bacterium             AKIFFVDKIIMVEGEIDAYFFEHYLQYLHNLPQGKGKE.....................
HEV2339438.1_Patescibacteria_group_bacterium            IKIFFSKKVVLIEGDSDEYFYRKFYDYYKEIKNKD....P...................
MBL9158386.1_Verrucomicrobiales_bacterium               VPLLRTKAVVFTENRDDRDYLKMFAVKHWGEEKTAR.......................
MGB8583703.1_Candidatus_Sulfotelmatobacter_sp.          SDALFYDRLICVEGKSDKQIIPLLLEKDGEIEPRELART....................
MBI3627515.1_Candidatus_Sungbacteria_bacterium          ISALDKKNNVIVEGPSDVFYLNAFKKTLNKN............................
_213105201.1_Candidatus_Protochlamydia_amoebophila      VYVNEPFAVVFVEGDKDKQFYSILAKRLNDTRKVE........................
CDRQKP010002909.1_Symbiodinium_sp.                      VKIETADNLIICEGISDIFYLRAFQKIFGVNNY..........................
JAUKPS010110286.1_Chlorellidium_tetrabotrys             IVGSDKLDNVVVEGPADYFYLTAFKQVVDGA............................
XP_004364654.1_Capsaspora_owczarzaki                    ....TVRPILVVDGETDVKFIRQFAPRLVE.............................
KJE91799.1_Capsaspora_owczarzaki                        ....IDRPVLLLDGKNDVEFYKKYVPNLSK.............................
XP_048575736.1_Nematostella_vectensis                   REIFFSRHIIWVEGESDKRFIEALLGLFDEGRGALFKALQVERAERSFSNED..SNRSL
A0A6P8J3J2_Actinia_tenebrosa                            REIFFSRHIIWVEGESDKRFIEALLGLFDEGQSDLWKVLTEHCTRTLLSREE.....SA
XP_022780642.1_Stylophora_pistillata                    REIFFARYVIWVEGESDKRFIEALLRLFDEGNMELFNALVEPGMEAKLSRG........
A0A9X0CFV3_Desmophyllum_pertusum                        REIFFARYIIWVEGESDKRFIEALLRLFDEGNLELFKALSQPGAATNTSGG........
A0A6S7HLM0_Paramuricea_clavata                          RELFFTRRIIWVEGDTDRRFCEAMLKLIDEGNDVLWSILMDPQDIDEEYRGAPAGSTGT
XP_046856289.1_Xenia_sp.                                RELFFTRSVIWVEGDTDQRFCETMLKFIDEGNPVLWSTLLDPNDCFYECRIINNTNSAG
A0A813MIX4_Adineta_steineri                             VRLGVHRKLVYLESLDDLNFLHGIIRRTESDLLRL........................
UJR19038.1_Adineta_vaga                                 VRFGVHPKLLCLESHDDYTFLHGIIHRAKPELLHF........................
JAXIUW010014154.1_Cephalothrix_simula                   NSILFASSVLFVERPADKFILELLIKEILDSDSPN........................
A0A8B8DK01_Crassostrea_virginica                        RTLLFATKVLTVEGPTDREVVQALFTFIKTEGIAN..............DISY......
XP_070564142.1_Ptychodera_flava                         SEIFFAEAVLWVEGADEIKFLESLSGVLEDLLVK...............NYT..GGSDS
ACQM01009845.1_Saccoglossus_kowalevskii                 RELFFSSFVLWVEGPDDISVLESMVIVLKEGLTKR..............NILHIGNCSI
XP_078582628.1_Branchiostoma_floridae_japonicum         ..ILKTKCILIVENESDADVLKAWIKKVQKIDIED........................
JALCYT010000124.1_Branchiostoma_belcheri                ..TLKTKCILIVENETDAEVLKAWIKKLHKIDIHD........................
KAJ3014353.1_Thoreauomyces_humboldtii                   MLLGARRNLLFVENDGDWTRLQLWMDRINSSRARAFAE.....................
KAJ3188706.1_Gaertneriomyces_sp._JEL0708                CLLSAKRHLLFMEDESDWNLLQMWMNRIDALRAAAFNN.....................
XP_002110806.1_Trichoplax_adhaerens                     VHLNLPFRLVFVEGTDDEMFYQMIMNKIIYKRLFMPSIPL...................
RDD37009.1_Trichoplax_sp._H2                            VYVNSPFRMVFVEAYDDKLFYQMIQLKLTNESMLEATYPL...................
A0A8W8L770_Magallana_gigas                              RTLLFSTKVLLVEGPIDKEVVQGIFTQYKCNLLER..............GVIG......
A0A8W8L770_Magallana_gigas                             α11       β17            α12
                                                                     ★    ★

E8PLM2_Thermus_scotoductus
                                                                                                                      400
E8PLM2_Thermus_scotoductus                              ..........................................................C
UJG41740.1_Candidatus_Heimdallarchaeum_aukensis         ...........................................ELNL...SST..NFLN
MFW9996705_Candidatus_Odinarchaeota_archaeon            ..........................................................T
MCH8915407.1_Thaumarchaeota_archaeon                    .......................................................NISD
MHA1972560.1_Candidatus_Hodarchaeales_archaeon          ..........................................................E
NHI95076.1_Candidatus_Lokiarchaeota_archaeon            ..........................................................P
MEM2144305.1_Candidatus_Jordarchaeaceae_archaeon        ..........................................................R
AHB41284.1_candidate_division_SR1_bacterium             .......................................................VITN
HEV2339438.1_Patescibacteria_group_bacterium            ..........................................................E
MBL9158386.1_Verrucomicrobiales_bacterium               .......................................................VWEG
MGB8583703.1_Candidatus_Sulfotelmatobacter_sp.          ..........................................................G
MBI3627515.1_Candidatus_Sungbacteria_bacterium          ..........................................................S
_213105201.1_Candidatus_Protochlamydia_amoebophila      ..........................................................S
CDRQKP010002909.1_Symbiodinium_sp.                      ..........................................................N
JAUKPS010110286.1_Chlorellidium_tetrabotrys             ..........................................................N
XP_004364654.1_Capsaspora_owczarzaki                    ..........................................................K
KJE91799.1_Capsaspora_owczarzaki                        ..........................................................H
XP_048575736.1_Nematostella_vectensis                   SPDWTPPRSRSPSPTPS.HRGNDAYVQKLLQDPAYERKYYTAEELGM...IQE..AVRS
A0A6P8J3J2_Actinia_tenebrosa                            TTDGRSP.SRSPSPRPF.PHTYDTFIATLFQDPAYERQYFTDEQLSL...VQE..AVRS
XP_022780642.1_Stylophora_pistillata                    .PD......CTGHSLKQVTKGLSLAVQNLLLDPAYKRVYYGKEVLSA...IQQ..GIRS
A0A9X0CFV3_Desmophyllum_pertusum                        .PDG....SLSEHSSSH.GAGRSLPVENLLQDPAYERKYYTREVLST...IQE..AVRS
A0A6S7HLM0_Paramuricea_clavata                          VEDMDDGSDSSTDEVDY.GKIGINPLDNLFQTSAYKRKYFRAKVLGD...CQE..LARS
XP_046856289.1_Xenia_sp.                                LP.MDDEIEDSTDEIDY.GAMGIDPLENLFQSRAYGRKYYTKEELMQ...CQE..LARS
A0A813MIX4_Adineta_steineri                             ..........................................................P
UJR19038.1_Adineta_vaga                                 ..........................................................P
JAXIUW010014154.1_Cephalothrix_simula                   .........................................DEDLQT........RVCG
A0A8B8DK01_Crassostrea_virginica                        .........................................EMDMKDKLPACDQLDFTT
XP_070564142.1_Ptychodera_flava                         ..........................................KMLET...VLD..VLKY
ACQM01009845.1_Saccoglossus_kowalevskii                 G........................................LHHFER...ALD..LMKR
XP_078582628.1_Branchiostoma_floridae_japonicum         ..........................................................K
JALCYT010000124.1_Branchiostoma_belcheri                ..........................................................K
KAJ3014353.1_Thoreauomyces_humboldtii                   ..........................................................R
KAJ3188706.1_Gaertneriomyces_sp._JEL0708                ..........................................................K
XP_002110806.1_Trichoplax_adhaerens                     ............................KFKSHGYSL.............RKHPPDECG
RDD37009.1_Trichoplax_sp._H2                            ............................LFMSPGVKRK............RQDSSDDSS
A0A8W8L770_Magallana_gigas                              .........................................MNELYK........DITT
A0A8W8L770_Magallana_gigas
```

```
                                                   β15            α10                              β16
E8PLM2_Thermus_scotoductus
                                                          410            420                  430
E8PLM2_Thermus_scotoductus                         IAVVDCGGKAGIELIVG......VCKALDIP..................FVVVHD
UJG41740.1_Candidatus_Heimdallarchaeum_aukensis    WDVISIDGTGDISKALS......IADFFDIK..................YVLICD
MFW9996705_Candidatus_Odinarchaeota_archaeon       ADIVNLHGKDEMTKAVK......VANHFSID..................WVVIAD
MCH8915407.1_Thaumarchaeota_archaeon               IEILNLEGKTNYPFVKN......FLNEFRIK..................SYFIGD
MHA1972560.1_Candidatus_Hodarchaeales_archaeon     INLVPGCG.DNIPAIAS......ILFGWGLD..................PYFILD
NHI95076.1_Candidatus_Lokiarchaeota_archaeon       YVIISRGGVDKNHYYEE.....SKLISEFIDS.................KIKVFSIID
MEM2144305.1_Candidatus_Jordarchaeaceae_archaeon   FTLLHLGGKGKILAFSE......TFEKVMLE............IVSRGFKIFALLD
AHB41284.1_candidate_division_SR1_bacterium        YEIININGKGSYKKRSA......FLKKFGIE..................SYFIGD
HEV2339438.1_Patescibacteria_group_bacterium       LEFLYISGKGSFELWKN......FLNKYKVK..................TYFIGD
MBL9158386.1_Verrucomicrobiales_bacterium          LSFLFTYQEPISADVKRLARQVKDLLNAPSLG...........DLAVGQQARFVVVGD
MGB8583703.1_Candidatus_Sulfotelmatobacter_sp.     FPILEGIGKGSTGSQTSILRH.EKLLSAIGRA.................DQPRMYLLD
MBI3627515.1_Candidatus_Sungbacteria_bacterium     VNFIFGGGCGNMPFVGT......ILHGWGGK..................VIYLYD
_213105201.1_Candidatus_Protochlamydia_amoebophila QMIFKVHGPGKQDNLSS.CGVIKSFIEKVTGQ.........TSEPLSPFVFGLID
CDRQKP010002909.1_Symbiodinium_sp.                 IKFVNGNG.DDISKIFF......ILEGTSNN..................GVRILD
JAUKPS010110286.1_Chlorellidium_tetrabotrys        INFVSGGGAGNMPKIGT......ILQGWGAR..................VIYLYD
XP_004364654.1_Capsaspora_owczarzaki               FTIVPSGSAGTDGSWNHRIA...FARALHLAGFKP..............PIFLKD
KJE91799.1_Capsaspora_owczarzaki                   FQLIS.GGTDTRHQLPTLTKL..LIETVNQP..................LFILRD
XP_048575736.1_Nematostella_vectensis              CKVLSVGGKMNLWKGAA......ICRDLHIP..................YAIICD
A0A6P8J3J2_Actinia_tenebrosa                       CKVVSIGGKKNLWKGAG......ICRDLRIP..................FAVICD
XP_022780642.1_Stylophora_pistillata               CAVLSINGKKNIYKAKA......ICHDLGIP..................HAVVCD
A0A9X0CFV3_Desmophyllum_pertusum                   CVVLSISGKKNVYKATA......ICNDLGIP..................HAVVCD
A0A6S7HLM0_Paramuricea_clavata                     CSVLPLSGKKNIDKATR......TCKDLGIP..................YAVICD
XP_046856289.1_Xenia_sp.                           CSILPLSGKKNMDKATK......ICKDLRIP..................FAVICD
A0A813MIX4_Adineta_steineri                        YTSIAKGGRPTSFQIEQ......LIVLFRKF............LPDKVKLHIFLLVD
UJR19038.1_Adineta_vaga                            FTRMARNGRTSPSQIKA......LISELRQL............LPKETTMDIFILVD
JAXIUW010014154.1_Cephalothrix_simula              MIVTELGDKREYKKASL......LVQMLHLQ..................SKYVFD
A0A8B8DK01_Crassostrea_virginica                   NQIIPLGSCENAKKVRS......FCKFIHLP..................CLCLLD
XP_070564142.1_Ptychodera_flava                    HTVLPLNGKDNLGKVG.......MCDQLQIS..................KVVICD
ACQM01009845.1_Saccoglossus_kowalevskii            ATILPLGGKFNISRGSQ......MCDDLGIA..................KAVVCD
XP_078582628.1_Branchiostoma_floridae_japonicum    IVVHSNSGRPTPEVVGK......LVGFLDRNLF.....HKKKSEQVVRESAKVFILTD
JALCYT010000124.1_Branchiostoma_belcheri           IVVHCNSTRPTPEVVGK......LVGFLDKNLF.....HQDEKCEKEQEKAKVFILTD
KAJ3014353.1_Thoreauomyces_humboldtii              VVVQSTSGRKPVYA.........YCEKLKLTTLVLSELAKVAGDDRAFEPLRVAISVD
KAJ3188706.1_Gaertneriomyces_sp._JEL0708           VVADWLGGKKAAHVYAD......VAKSMGNW...MRE.......STKTQKLKVAVVVD
XP_002110806.1_Trichoplax_adhaerens                NITTEDSSRQIIERLVK......FCVDPKVA..........DECDDYSLKGFIFGLVD
RDD37009.1_Trichoplax_sp._H2                       SKMIENSCRALVENIVE......TCMNIENE............DLSLREFVFGLID
A0A8W8L770_Magallana_gigas                         SQIIPVGGCENTKKIQS......FCYFINLP..................CLCLWD
A0A8W8L770_Magallana_gigas
                                                     β18               α13                                   β19
                                                                                                                ★

                                                                                                              α11
E8PLM2_Thermus_scotoductus                                          TT
                                                                   440          450
E8PLM2_Thermus_scotoductus                         EDVW.........PIDERADEETRRKQEQEN..........................
UJG41740.1_Candidatus_Heimdallarchaeum_aukensis    FDAF.........LKPKNSLTNSYISKILKQ..RCNINLDEIFSSKTTNIQE......
MFW9996705_Candidatus_Odinarchaeota_archaeon       FDAW.......KPIGGKSLSESRIGGILSQ..DTSFDSSKYSFLKKNDPYE.......
MCH8915407.1_Thaumarchaeota_archaeon               LDNIFTLP.........................NPILGPNLKLRFGGGNT........
MHA1972560.1_Candidatus_Hodarchaeales_archaeon     SDKQ..............RIVNKLKKKLDI............................
NHI95076.1_Candidatus_Lokiarchaeota_archaeon       RDFR............SVEESDYYIENSKKH...........................
MEM2144305.1_Candidatus_Jordarchaeaceae_archaeon   KDKE......................................................
AHB41284.1_candidate_division_SR1_bacterium        WDNIVDYG......FISQQDLNFYYKQAKTFYHTH.......KKDKETTDR.......
HEV2339438.1_Patescibacteria_group_bacterium       WDNVVSFN......LLSRSQLNKFNKAYLNSVYTN...IAVNVLSKKGSTDRVNMLSSL
MBL9158386.1_Verrucomicrobiales_bacterium          RDYR............SEASSEAARNDLESK...........................
MGB8583703.1_Candidatus_Sulfotelmatobacter_sp.     GDRK.............DDERTVIQGTKHP............................
MBI3627515.1_Candidatus_Sungbacteria_bacterium     NDQG.............KKDGEKNLKDNWLV...........................
_213105201.1_Candidatus_Protochlamydia_amoebophila GDNH......................................................
CDRQKP010002909.1_Symbiodinium_sp.                 TDEN..............KIKKKMKEFIKI............................
JAUKPS010110286.1_Chlorellidium_tetrabotrys        NDQA............FKDAQRSIKKEWLTI...........................
XP_004364654.1_Capsaspora_owczarzaki               SEMM...........ASSSRLDNLQRAYLAD...........................
KJE91799.1_Capsaspora_owczarzaki                   SELRACWP..ESLLAWGEYCLSNVQKEVRP............................
XP_048575736.1_Nematostella_vectensis              LDAI...........VPNSRENSIQSQFERA..EGDWYSASIPHAKAKLVDE..EEAPA
A0A6P8J3J2_Actinia_tenebrosa                       LDAI...........VPNSRENSILSQFEKC..DGNWEAAKIPHAKTRLVDE..EESPA
XP_022780642.1_Stylophora_pistillata               LDVI...........IPNSKENSVQSQFNKC..NGVWANACIPNAKTKLLDE..DQCPA
A0A9X0CFV3_Desmophyllum_pertusum                   LDVI...........IPNSKENSVQSQFDSC..RGLWTSAPIPHAKTKLLNE..DQCPA
A0A6S7HLM0_Paramuricea_clavata                     LDAV...........LPNSKKNSIMSQFMKC..VGNWSLGYV..EKKLLFDS..EQSPA
XP_046856289.1_Xenia_sp.                           LDAV...........LPNSRKNSIMSQFLGS..VGNWSLAPA..DKKLLFDG..EQSPA
A0A813MIX4_Adineta_steineri                        ADLR............SEDILEEEKKSYINL...........................
UJR19038.1_Adineta_vaga                            ADLR............SKSVVDEEEQSYDEL...........................
JAXIUW010014154.1_Cephalothrix_simula              QDEE....NDKHLFNIPDKPEEEVEKYLKL............................
A0A8B8DK01_Crassostrea_virginica                   LDKF............VSVKKEDNTIENFKSI..........................
XP_070564142.1_Ptychodera_flava                    LDAV..........LPADSDLKTYRKQNDNS.........QSLTSTVEE.RKAVSA
ACQM01009845.1_Saccoglossus_kowalevskii            RDAVISTLGEKLTKTAAASSLNATEKKSLEQ...........................
XP_078582628.1_Branchiostoma_floridae_japonicum    RDYK............AKMALEKEQRLLDEN...........................
JALCYT010000124.1_Branchiostoma_belcheri           RDYK............ADMALEKEQRILREN...........................
KAJ3014353.1_Thoreauomyces_humboldtii              GDYI............PAPVLQHEQKQFRIL...........................
KAJ3188706.1_Gaertneriomyces_sp._JEL0708           GDYK........PIALQRKDRDSFLEEAREA...........................
XP_002110806.1_Trichoplax_adhaerens                NDNK............GESK......................................
RDD37009.1_Trichoplax_sp._H2                       NDNK............EEPK......................................
A0A8W8L770_Magallana_gigas                         LDKV...........VKFDKQTCKIQDFLEI...........................
A0A8W8L770_Magallana_gigas
                                                   η11             β20   TTT      β21
                                                    ★
```

```
                                                                                                              η5
E8PLM2_Thermus_scotoductus                         .........................                                  
                                                                              460                        470
E8PLM2_Thermus_scotoductus                         .........................KAEQEKNQRIQACAG...................AERV
UJG41740.1_Candidatus_Heimdallarchaeum_aukensis    ............................RKKEIISFAK.......................NYNI
MFW9996705_Candidatus_Odinarchaeota_archaeon       ...........................QNVRNGLHKLER......................SHGI
MCH8915407.1_Thaumarchaeota_archaeon               ............................TNKQIIQNIKANQQTEWSQIRVEIFLRYFYNV
MHA1972560.1_Candidatus_Hodarchaeales_archaeon     ............................SEDVIIKILDE.......................
NHI95076.1_Candidatus_Lokiarchaeota_archaeon       ..............................................................QVDV
MEM2144305.1_Candidatus_Jordarchaeaceae_archaeon   ...............................................SFAA....KSDFC
AHB41284.1_candidate_division_SR1_bacterium        ............................YYAKLVNAVKYLYPNKYSSILEKIDFLYTKNV
HEV2339438.1_Patescibacteria_group_bacterium       SSYISHPTKQGFKKLKEMRGYMLVRYIPSEDLVNFIKTDAST.YSTLQTSIADLYKKQI
MBL9158386.1_Verrucomicrobiales_bacterium          ............................AKSDDFKL.......................DLVC
MGB8583703.1_Candidatus_Sulfotelmatobacter_sp.     ...........................................................TTLSSLTI
MBI3627515.1_Candidatus_Sungbacteria_bacterium     ............................SKDLILSVLGT.......................
_213105201.1_Candidatus_Protochlamydia_amoebophila ..........................................................NHVGTARNL
CDRQKP010002909.1_Symbiodinium_sp.                 ............................QSNNILEISEE..K...................
JAUKPS010110286.1_Chlorellidium_tetrabotrys        ............................SKDCLLAL...NV....................
XP_004364654.1_Capsaspora_owczarzaki               ...........................................................FAGAAAGV
KJE91799.1_Capsaspora_owczarzaki                   ..............................................................RCQI
XP_048575736.1_Nematostella_vectensis              SKAIDETEHAQLAKLR.......SLRTVGEVIRFYEQ...................NHYI
A0A6P8J3J2_Actinia_tenebrosa                       SKAIDENDRDRIAKLK.......SLRYVGDVMRFYDK...................ELYF
XP_022780642.1_Stylophora_pistillata               SRCVRDSSPGLIPKLR.......ACRTVAEVMNFYDK...................MQHI
A0A9X0CFV3_Desmophyllum_pertusum                   SKCVQDSNPGLIPKLR.......ACRTVGEVMKFYDE...................TQHI
A0A6S7HLM0_Paramuricea_clavata                     STCIEDPSLK..AMLR.......KCHNAHEIMEFYET...................MERI
XP_046856289.1_Xenia_sp.                           FTCLEDTSAQ..EELR.......KCRNALDVMEFYEK...................NERI
A0A813MIX4_Adineta_steineri                        ............................QENENLNV.......................RIHY
UJR19038.1_Adineta_vaga                            ............................QKDTTLNV.......................KVYY
JAXIUW010014154.1_Cephalothrix_simula              ..............................................................KRSI
A0A8B8DK01_Crassostrea_virginica                   ......PDKQYEEKLT.......DDFTTKHISSFFSNGFDEFSKFLKT.....EMDT
XP_070564142.1_Ptychodera_flava                    AKCLAKDIKGGL..........ADCKTVKDILDFYQ....................EHRV
ACQM01009845.1_Saccoglossus_kowalevskii            ......................TCTTDKDVIDFFE.....................KHHI
XP_078582628.1_Branchiostoma_floridae_japonicum    ............................TKKYNGRI.......................KVEW
JALCYT010000124.1_Branchiostoma_belcheri           ............................TKRYV.RI.......................KVEW
KAJ3014353.1_Thoreauomyces_humboldtii              ............................LKKECGDSVDAD...................SCLW
KAJ3188706.1_Gaertneriomyces_sp._JEL0708           ....................................RCLEPK..................GLFW
XP_002110806.1_Trichoplax_adhaerens                ..............................................................INNI
RDD37009.1_Trichoplax_sp._H2                       ..............................................................LYNI
A0A8W8L770_Magallana_gigas                         ......GNVNK.DKLK.......AKYFDKELSYLI..GCDEDFETISTS...LEQRKM
A0A8W8L770_Magallana_gigas                               TT                                α15

                                                   β17            α12                                        η6
E8PLM2_Thermus_scotoductus                         
                                                            480                        490
E8PLM2_Thermus_scotoductus                         FVV.....QPSLE...........AALGIGRNAS.......................
UJG41740.1_Candidatus_Heimdallarchaeum_aukensis    FSW.....ENDLE...........GMIKKTV..........................
MFW9996705_Candidatus_Odinarchaeota_archaeon       FTWKCTGDKGDLE...........SAIQETL..........................
MCH8915407.1_Thaumarchaeota_archaeon               FLLR....EGELEDYIGRSQGKLENVIIFCRDHF.......................
MHA1972560.1_Candidatus_Hodarchaeales_archaeon     ........KGTVED..VFSEDDFKKFVIEDEDA...DLTGGFIS.............
NHI95076.1_Candidatus_Lokiarchaeota_archaeon       FILD....KKEIENY.LIVPQLVMRCINESRI......KRGLKQL............
MEM2144305.1_Candidatus_Jordarchaeaceae_archaeon   FTWP....RACIENFLLLDSGAIYEALKVTVG......EVSLQDK............
AHB41284.1_candidate_division_SR1_bacterium        FILK....KGDIETYLGIQSKGLEETIAFCHNFFD......................
HEV2339438.1_Patescibacteria_group_bacterium       YILQ....EGELEEYLGTTGKGLDVVIAFCKNDF.......................
MBL9158386.1_Verrucomicrobiales_bacterium          SIWK....RNEIENY.LLDPSALEEAVVSTLR......DSSLEQKARDVVKA.....
MGB8583703.1_Candidatus_Sulfotelmatobacter_sp.     KFLP....RPEIENY.LLAPEAIAAALHEELILKGEPGEVSLTEVSQ..........
MBI3627515.1_Candidatus_Sungbacteria_bacterium     ........AGSIED..IFSPSDFKQFVLNDVDK...TYTGTNAE.............
_213105201.1_Candidatus_Protochlamydia_amoebophila KALK....RYSIENY.ILDPIHLFLCLNFIRENF..STDSDLQPHVKDCVEK.FEALVS
CDRQKP010002909.1_Symbiodinium_sp.                 ........NKIIED..LFEEKDFKRFVIGNKEV...DLLSNRKEF............
JAUKPS010110286.1_Chlorellidium_tetrabotrys        ........DGAIED..IFSKSEFASILDVAPGE...ILEKNSL..............
XP_004364654.1_Capsaspora_owczarzaki               YFTR....LPCIESY.LYLHLLLASPANAEKV.........LKEKLSPML.......
KJE91799.1_Capsaspora_owczarzaki                   FFTS....LPCIESY.LFVHCALTDSSFADDQ......AAQVKQKLCSSKSV.....
XP_048575736.1_Nematostella_vectensis              FAWWP.M.GGEIE...........DAVRLTK..........................
A0A6P8J3J2_Actinia_tenebrosa                       FTWRL.M.GGEIE...........DAVRLTR..........................
XP_022780642.1_Stylophora_pistillata               FTWHV.D.GGEIE...........DAIRLTK..........................
A0A9X0CFV3_Desmophyllum_pertusum                   FTWRV.D.GGEIE...........DAIRLTK..........................
A0A6S7HLM0_Paramuricea_clavata                     FAWRV.G.NGEIE...........DLVKVTK..........................
XP_046856289.1_Xenia_sp.                           FAWRV.G.NGEIE...........DVVKVTK..........................
A0A813MIX4_Adineta_steineri                        HCWA....AREWENW.LLSNGDLLYEMLFSDHLSD...AEEIK..............
UJR19038.1_Adineta_vaga                            HCWA....AREWENW.LLYNEDLLRDILFGNKLPS...IKEIHALRIEIFGS.....
JAXIUW010014154.1_Cephalothrix_simula              FLWK....QGNLEKM.LFKAIYDDKVILPDMKIL.......................
A0A8B8DK01_Crassostrea_virginica                   FVWK.....GTLEDA.ILSSD.....................................
XP_070564142.1_Ptychodera_flava                    FAWSV.G.DGNIE...........DIIQIAQ..........................
ACQM01009845.1_Saccoglossus_kowalevskii            YSWS....DGSIENFFVLDP....TVLEHKGK.........................
XP_078582628.1_Branchiostoma_floridae_japonicum    YCYK....KNEQENY.FLSAAALKSYLEEKGKS.....TENLE..............
JALCYT010000124.1_Branchiostoma_belcheri           YCYK....KNEQENN.FLSATALKSYLEEKGGK.....TESMEKI............
KAJ3014353.1_Thoreauomyces_humboldtii              QTWK....QVESENY.LLDKDLIVRFLESGTIKGK.AGGLAPEELWNSVMAT.....
KAJ3188706.1_Gaertneriomyces_sp._JEL0708           HSWS....CVEIENY.LLDEEAIFRVLFPEKKQM....SEDFRQKFRTLLEY.....
XP_002110806.1_Trichoplax_adhaerens                TCLKE...RYSLENY.IYDPVYLYYNLLRRGEVL....VSQIKEEINSKLCLQNHPDHL
RDD37009.1_Trichoplax_sp._H2                       RCLN....RYSMENY.LYDPVYIYYYLRKCGK......AEEIKTQINVNLQEKQHPDCL
A0A8W8L770_Magallana_gigas                         FVWR....HGTVEDA.ILSSR.....................................
A0A8W8L770_Magallana_gigas                         β22  T....T
```

```
                                                          α13
*E8PLM2_Thermus_scotoductus*                                                    T
                                                        500

E8PLM2_Thermus_scotoductus                         .............DKPYRIAE.......ILKTVDV........................
UJG41740.1_Candidatus_Heimdallarchaeum_aukensis    ...........................................................
MFW9996705_Candidatus_Odinarchaeota_archaeon       ...........................................................
MCH8915407.1_Thaumarchaeota_archaeon               ...........................................................
MHA1972560.1_Candidatus_Hodarchaeales_archaeon     ............................LIKDN..........................
NHI95076.1_Candidatus_Lokiarchaeota_archaeon       ..........SQEDVNNMLNE.......ICESMKE......STIDKYSQSLLAFNRKID
MEM2144305.1_Candidatus_Jordarchaeaceae_archaeon   .......GIKTEKDIDNLIEE.......IIKNPEV.......LKEEIKQRAEEHLKFHI
AHB41284.1_candidate_division_SR1_bacterium        ...........................................................
HEV2339438.1_Patescibacteria_group_bacterium       ...........................................................
MBL9158386.1_Verrucomicrobiales_bacterium          .................SLSK.......NLVSLKE......DARLRIAGKLQHENHEFR
MGB8583703.1_Candidatus_Sulfotelmatobacter_sp.     .................RLNE.......LLNQENK........................
MBI3627515.1_Candidatus_Sungbacteria_bacterium     ............................YVKKAK.........................
_213105201.1_Candidatus_Protochlamydia_amoebophila SLPLVEESIEKQMSDPNFCQQ.......ILNVFYT..........RLRKECEKALSCRE
CDRQKP010002909.1_Symbiodinium_sp.                 ............................FLKSNIG........................
JAUKPS010110286.1_Chlorellidium_tetrabotrys        ............................YLKSNK.........................
XP_004364654.1_Capsaspora_owczarzaki               .................ALEESRDYFVEIYKQHSS......KLRPSLSNESV.......
KJE91799.1_Capsaspora_owczarzaki                   .TRFI..NYFTEQNFQGAINE......................HQRASNEEQPHP....
XP_048575736.1_Nematostella_vectensis              ...........................................................
A0A6P8J3J2_Actinia_tenebrosa                       ...........................................................
XP_022780642.1_Stylophora_pistillata               ...........................................................
A0A9X0CFV3_Desmophyllum_pertusum                   ...........................................................
A0A6S7HLM0_Paramuricea_clavata                     ...........................................................
XP_046856289.1_Xenia_sp.                           ...........................................................
A0A813MIX4_Adineta_steineri                        .................QLQD.......IMKRVWSTDLVFPTNRDQF............
UJR19038.1_Adineta_vaga                            ......STLQPNPAMHSRLEKSESAV..VASTIGTSVSQLENSKDKF............
JAXIUW010014154.1_Cephalothrix_simula              ...LI...........YSIQN.......IQGNCGPVFLS....................
A0A8B8DK01_Crassostrea_virginica                   ................EHEND.......IAKSLGLTFTS....................
XP_070564142.1_Ptychodera_flava                    ...........................................................
ACQM01009845.1_Saccoglossus_kowalevskii            ...........................................................
XP_078582628.1_Branchiostoma_floridae_japonicum    ................TKLND.......LVDKHCQAF....TIREKYMNSVRYVTREKG
JALCYT010000124.1_Branchiostoma_belcheri           ................KKLDD.......LLKKHCRAF....SIREKYMNSVRNILKEKG
KAJ3014353.1_Thoreauomyces_humboldtii              .................ELAE.......VERRFAA.........ESASGASIKGDDNLA
KAJ3188706.1_Gaertneriomyces_sp._JEL0708           .................QLQA.......IINRFAC..........EYVEAFRAGTRKLT
XP_002110806.1_Trichoplax_adhaerens                SDFL.....EGTETKIDILQA.......IIDNVFD..........KLAKTIERFIRLSE
RDD37009.1_Trichoplax_sp._H2                       SDLLME...KNVTNLTKVLQQ.......IVNIMSK..........NLFDIVEKFIKLKD
A0A8W8L770_Magallana_gigas                         ................NRYEE.......ICSSINC........................
*A0A8W8L770_Magallana_gigas*
                                                          α18


*E8PLM2_Thermus_scotoductus*                       .........................................T

E8PLM2_Thermus_scotoductus                         .........................................G.....Q...........
UJG41740.1_Candidatus_Heimdallarchaeum_aukensis    ........................TKFKKDDW...........................
MFW9996705_Candidatus_Odinarchaeota_archaeon       ........................PHFRKKEW...........................
MCH8915407.1_Thaumarchaeota_archaeon               ........................SNWKNT.............................
MHA1972560.1_Candidatus_Hodarchaeales_archaeon     ......................GGKELLAKKFLEKVDSKEI..................
NHI95076.1_Candidatus_Lokiarchaeota_archaeon       KK..CDLTLTNPIA..........RDIVAKKWTD.LN......................
MEM2144305.1_Candidatus_Jordarchaeaceae_archaeon   TGDWEDLEGLKKVS..........SEILDKRLAR..............FEE........
AHB41284.1_candidate_division_SR1_bacterium        ........................VRLQNKNLAEHRK......................
HEV2339438.1_Patescibacteria_group_bacterium       ........................ENWIKDQA...........................
MBL9158386.1_Verrucomicrobiales_bacterium          G....DYVKTTKEA..........SRILDEEW..G........................
MGB8583703.1_Candidatus_Sulfotelmatobacter_sp.     ..............PP.....VKGSRVLEKLY...........................
MBI3627515.1_Candidatus_Sungbacteria_bacterium     ......................LDKVLLAKKFLETCQNGT...................
_213105201.1_Candidatus_Protochlamydia_amoebophila QN.FADLATLIQEQIDVCEAILENKKISGKLFDTIEQMIKKPGKAIRFFKEFLLGLQKE
CDRQKP010002909.1_Symbiodinium_sp.                 ......................KSKALMSKIFLQKVDNGTL..................
JAUKPS010110286.1_Chlorellidium_tetrabotrys        ......................RDKVLPARHFLERMRSDT...................
XP_004364654.1_Capsaspora_owczarzaki               ........................QEYATRLWND.........................
KJE91799.1_Capsaspora_owczarzaki                   ........................VLEACRRWEAGKTFLT...................
XP_048575736.1_Nematostella_vectensis              ........................AQFGKKLW...........................
A0A6P8J3J2_Actinia_tenebrosa                       ........................AQFGKKLW...........................
XP_022780642.1_Stylophora_pistillata               ........................SQFGKKLW...........................
A0A9X0CFV3_Desmophyllum_pertusum                   ........................SQFGKKLW...........................
A0A6S7HLM0_Paramuricea_clavata                     ........................RTFSKKRW...........................
XP_046856289.1_Xenia_sp.                           ........................PKFSKKRW...........................
A0A813MIX4_Adineta_steineri                        ........................NEWFEKKLDHHFKLLK...................
UJR19038.1_Adineta_vaga                            ........................CEWFAKKLEHHFSVLL...................
JAXIUW010014154.1_Cephalothrix_simula              ........................QDSREKIFGNS........................
A0A8B8DK01_Crassostrea_virginica                   ......................ENPKERAKQLKRKLK......................
XP_070564142.1_Ptychodera_flava                    ........................KSQKRIKL...........................
ACQM01009845.1_Saccoglossus_kowalevskii            ......................KRKGPLKKSI...........................
XP_078582628.1_Branchiostoma_floridae_japonicum    GK......AMLKEPTF.....HQKTEEEPKGWLEG........................
JALCYT010000124.1_Branchiostoma_belcheri           GK......GQLKDPEF.....HQETEDETKTWLA.........................
KAJ3014353.1_Thoreauomyces_humboldtii              KADRFNQSELMPSL..........MDYMTQRWQEARR......................
KAJ3188706.1_Gaertneriomyces_sp._JEL0708           PKDIAHMFPL.........GL...LGEMNRILDAAKE......................
XP_002110806.1_Trichoplax_adhaerens                SKDFKDLVSLKQHLNK....LKRMNDQLNEYQKIGEEIKE.......DLKE.LKRIYKL
RDD37009.1_Trichoplax_sp._H2                       ..DFQEAINSLKTLHKD...VVSKSGILNKESIITID......................
A0A8W8L770_Magallana_gigas                         ........................PSLTNKSLKKKLK......................
*A0A8W8L770_Magallana_gigas*
```

```
E8PLM2_Thermus_scotoductus                                                                 .........
                                                                                    510
E8PLM2_Thermus_scotoductus                             .........................................PPDALRPFV.........
UJG41740.1_Candidatus_Heimdallarchaeum_aukensis        .........................................KDMTFEECF.........
MFW9996705_Candidatus_Odinarchaeota_archaeon           .........................................KEKSYHETI.........
MCH8915407.1_Thaumarchaeota_archaeon                   .........................................NTNYLQEFN.........
MHA1972560.1_Candidatus_Hodarchaeales_archaeon         ....................................KKNSLSKQTIENIE.........
NHI95076.1_Candidatus_Lokiarchaeota_archaeon           .........................................GKISCCDGS.........
MEM2144305.1_Candidatus_Jordarchaeaceae_archaeon       .................................QYEKLNNEIKQVVNDKE.........
AHB41284.1_candidate_division_SR1_bacterium            ...............................................ELD.........
HEV2339438.1_Patescibacteria_group_bacterium           .........................................NTAKIDEIT.........
MBL9158386.1_Verrucomicrobiales_bacterium              .........................................DGLALCDAK.........
MGB8583703.1_Candidatus_Sulfotelmatobacter_sp.         .........................................DSFSLRYLK.........
MBI3627515.1_Candidatus_Sungbacteria_bacterium         .....................................SISLDKTTMDNIN.........
_213105201.1_Candidatus_Protochlamydia_amoebophila     FEQTIEKKNCDIFNIAGNLCFDEKNANLFKKNPDKEAVVKKLKSFFEDRK.........
CDRQKP010002909.1_Symbiodinium_sp.                     ....................................KKDNFNKTTIDNFQ.........
JAUKPS010110286.1_Chlorellidium_tetrabotrys           .....................................PPQLSAQTVVVIK.........
XP_004364654.1_Capsaspora_owczarzaki                   .........................................RLQALRNAK.........
KJE91799.1_Capsaspora_owczarzaki                       .........................................APDSPRDAR.........
XP_048575736.1_Nematostella_vectensis                  .........................................PDMLSEEIK.........
A0A6P8J3J2_Actinia_tenebrosa                           .........................................PDMLSAELK.........
XP_022780642.1_Stylophora_pistillata                   .........................................PDLSFEEVK.........
A0A9X0CFV3_Desmophyllum_pertusum                       .........................................PDLSFGDIK.........
A0A6S7HLM0_Paramuricea_clavata                         .........................................ADFTSDAFQ.........
XP_046856289.1_Xenia_sp.                               .........................................ADFTAENFQ.........
A0A813MIX4_Adineta_steineri                            .........................................LELSLSQMG.......YV
UJR19038.1_Adineta_vaga                                .........................................TKLTLSEMH.......GT
JAXIUW010014154.1_Cephalothrix_simula                  .........................................CDNNMTDDE.........
A0A8B8DK01_Crassostrea_virginica                       .........................................KRLTDEERQ.........
XP_070564142.1_Ptychodera_flava                        .........................................ATFSEQDMD.........
ACQM01009845.1_Saccoglossus_kowalevskii                .........................................VEYSADELD.........
XP_078582628.1_Branchiostoma_floridae_japonicum        .........................................QRSSFTDAK.........
JALCYT010000124.1_Branchiostoma_belcheri               .........................................QPSSFTDAK.........
KAJ3014353.1_Thoreauomyces_humboldtii                  .........................................APATIVNAK.........
KAJ3188706.1_Gaertneriomyces_sp._JEL0708               .........................................NPRILVDAK.........
XP_002110806.1_Trichoplax_adhaerens                    NSNQFSKQSNGQGNIQNYIQNNSQNTDQVCVQHNDQGINQSDKQSYIDGQLMRKIKITL
RDD37009.1_Trichoplax_sp._H2                           ........................NFETV............CKFVYNEGKLIDYVKKIR
A0A8W8L770_Magallana_gigas                             .........................................ERLDEKERR.........
A0A8W8L770_Magallana_gigas                                                                      .........

                                                            α15
E8PLM2_Thermus_scotoductus                             ..............................
                                                              ★520
E8PLM2_Thermus_scotoductus                             ..............................EAIRQVTRPMEE.................
UJG41740.1_Candidatus_Heimdallarchaeum_aukensis        ..............................NLVIKLIKNKNT.................
MFW9996705_Candidatus_Odinarchaeota_archaeon           ..............................ELAKELLNKDNA.................
MCH8915407.1_Thaumarchaeota_archaeon                   ..............................FIFTNILNDH...................
MHA1972560.1_Candidatus_Hodarchaeales_archaeon         ..............................GIFEKISDLLNELNDS.............
NHI95076.1_Candidatus_Lokiarchaeota_archaeon           ..............................DILNEINKKMSP.................
MEM2144305.1_Candidatus_Jordarchaeaceae_archaeon       ..............................KALVELNGKIILGKISSRFNVRRDYLAR.
AHB41284.1_candidate_division_SR1_bacterium            ..............................HILFTIFSQ....................
HEV2339438.1_Patescibacteria_group_bacterium           ..............................KIFDSITA.....................
MBL9158386.1_Verrucomicrobiales_bacterium              ..............................KLLSGIRADLQSARIPARLN.........
MGB8583703.1_Candidatus_Sulfotelmatobacter_sp.         ..............................ESSGVLIARHIT.................
MBI3627515.1_Candidatus_Sungbacteria_bacterium         ..............................KLIENIEGKF...................
_213105201.1_Candidatus_Protochlamydia_amoebophila     ESADKKYEGDWA..................KLVNCLGQRDTKIMDESDII...DFDSKE
CDRQKP010002909.1_Symbiodinium_sp.                     ..............................KILEKLK......................
JAUKPS010110286.1_Chlorellidium_tetrabotrys           ..............................SLFALLDEKFKDYKS..............
XP_004364654.1_Capsaspora_owczarzaki                   .......DHDFG...............FWRGLVCLLKGHALYKTS..............
KJE91799.1_Capsaspora_owczarzaki                       ...........................YWRTYVNLVGSHEIFGSKDT............
XP_048575736.1_Nematostella_vectensis                  ..............................SFILSLIDPRKLLIKNPK...........
A0A6P8J3J2_Actinia_tenebrosa                           ..............................ELIISLLDPRKLLLKDPK...........
XP_022780642.1_Stylophora_pistillata                   ..............................ELVVGLLQPCKLQQGSSQ...........
A0A9X0CFV3_Desmophyllum_pertusum                       ..............................ELVVCLLQPPKLRERNPR...........
A0A6S7HLM0_Paramuricea_clavata                         ..............................ELIQFMLVPSVEDTPG.............
XP_046856289.1_Xenia_sp.                               ..............................DLMQMMLSPASSGYQG.............
A0A813MIX4_Adineta_steineri                            KTPD.......................................................
UJR19038.1_Adineta_vaga                                ETDERKNAKQLAKQAGEDFLLKAGVTD...DILQNFGSFRNAIILSVGFQVNQKENPKK
JAXIUW010014154.1_Cephalothrix_simula                  ............L..............C..GDLGKVLRRSE..................
A0A8B8DK01_Crassostrea_virginica                       ..............................KMCTQLMK.....................
XP_070564142.1_Ptychodera_flava                        ..............................GVCQTLINSNHD.................
ACQM01009845.1_Saccoglossus_kowalevskii                ..............................GILTNILSSSME.................
XP_078582628.1_Branchiostoma_floridae_japonicum        ..............................EVLKELGIEEDP.................
JALCYT010000124.1_Branchiostoma_belcheri               ..............................GVLKELGVEEES.................
KAJ3014353.1_Thoreauomyces_humboldtii                  ..............................SALKALGVAPDL.................
KAJ3188706.1_Gaertneriomyces_sp._JEL0708               ..............................EVLKELNIARHD.................
XP_002110806.1_Trichoplax_adhaerens                    ETENTKYDT................LT...ERLRNLCKYTDEITDAIKRL.KSNQFLRN
RDD37009.1_Trichoplax_sp._H2                           QRTNCK........................GLLKELIDYTTEIVEATNRLIKKKNLIKK
A0A8W8L770_Magallana_gigas                             ..............................TFYSQLME.....................
A0A8W8L770_Magallana_gigas                             ..............................
                                                            α21
```

*E8PLM2_Thermus_scotoductus*

```
E8PLM2_Thermus_scotoductus                              ...........................................................
UJG41740.1_Candidatus_Heimdallarchaeum_aukensis         ...........................................................
MFW9996705_Candidatus_Odinarchaeota_archaeon            ...........................................................
MCH8915407.1_Thaumarchaeota_archaeon                    ...........................................................
MHA1972560.1_Candidatus_Hodarchaeales_archaeon          ...........................................................
NHI95076.1_Candidatus_Lokiarchaeota_archaeon            ...........................................................
MEM2144305.1_Candidatus_Jordarchaeaceae_archaeon        ...........................................................
AHB41284.1_candidate_division_SR1_bacterium             ...........................................................
HEV2339438.1_Patescibacteria_group_bacterium            ...........................................................
MBL9158386.1_Verrucomicrobiales_bacterium               ...........................................................
MGB8583703.1_Candidatus_Sulfotelmatobacter_sp.          ...........................................................
MBI3627515.1_Candidatus_Sungbacteria_bacterium          ...........................................................
_213105201.1_Candidatus_Protochlamydia_amoebophila      SENVQL........ILKNQIINEH..ELQYP..................KFLLYFRGHN
CDRQKP010002909.1_Symbiodinium_sp.                      ...........................................................
JAUKPS010110286.1_Chlorellidium_tetrabotrys             ...........................................................
XP_004364654.1_Capsaspora_owczarzaki                    ...........................................................
KJE91799.1_Capsaspora_owczarzaki                        ...........................................................
XP_048575736.1_Nematostella_vectensis                   ...........................................................
A0A6P8J3J2_Actinia_tenebrosa                            ...........................................................
XP_022780642.1_Stylophora_pistillata                    ...........................................................
A0A9X0CFV3_Desmophyllum_pertusum                        ...........................................................
A0A6S7HLM0_Paramuricea_clavata                          ...........................................................
XP_046856289.1_Xenia_sp.                                ...........................................................
A0A813MIX4_Adineta_steineri                             ...........................................................
UJR19038.1_Adineta_vaga                                 HRSKVAADKPMDEEKCREKWLAER..QLQQPIVCGELATRKELTKWIDAKLFFHLLTHD
JAXIUW010014154.1_Cephalothrix_simula                   ...........................................................
A0A8B8DK01_Crassostrea_virginica                        ...........................................................
XP_070564142.1_Ptychodera_flava                         ...........................................................
ACQM01009845.1_Saccoglossus_kowalevskii                 ...........................................................
XP_078582628.1_Branchiostoma_floridae_japonicum         ...........................QSYT..................QLIHHFP...
JALCYT010000124.1_Branchiostoma_belcheri                ...........................QSYI..................RIIHHFL...
KAJ3014353.1_Thoreauomyces_humboldtii                   .............................YP..................R.........
KAJ3188706.1_Gaertneriomyces_sp._JEL0708                .............................YSS...........................
XP_002110806.1_Trichoplax_adhaerens                     GDNSKLDFS.......KSDISLESKVTVSYP..................IVILKMRGHD
RDD37009.1_Trichoplax_sp._H2                            QKGSEEAYEL...IKCTEDVRFENNITLSYP..................KVILKMRGHD
A0A8W8L770_Magallana_gigas                              ...........................................................
```

*A0A8W8L770_Magallana_gigas*

*E8PLM2_Thermus_scotoductus*

```
E8PLM2_Thermus_scotoductus                              ...........................................................
UJG41740.1_Candidatus_Heimdallarchaeum_aukensis         ...............................................EFLAFLEFLGNL
MFW9996705_Candidatus_Odinarchaeota_archaeon            ...............................................CIKDLISFLTQK
MCH8915407.1_Thaumarchaeota_archaeon                    ...........................................................
MHA1972560.1_Candidatus_Hodarchaeales_archaeon          ...........................................................
NHI95076.1_Candidatus_Lokiarchaeota_archaeon            .........................NMI.VINIGQLLSNISKEEISNDLIEIFKKLQ..
MEM2144305.1_Candidatus_Jordarchaeaceae_archaeon        ...........................T.IAD.....KLRILNRLPRQITDLIEQIEKV
AHB41284.1_candidate_division_SR1_bacterium             ...........................................................
HEV2339438.1_Patescibacteria_group_bacterium            ...........................................................
MBL9158386.1_Verrucomicrobiales_bacterium               ..................................EADVVKQMSEVPKDVRIVLQRLQQL
MGB8583703.1_Candidatus_Sulfotelmatobacter_sp.          ........................A......................SMQPIISELTDL
MBI3627515.1_Candidatus_Sungbacteria_bacterium          ...........................................................
_213105201.1_Candidatus_Protochlamydia_amoebophila      .........LQTCYEKILKI....PHEL.IKDFAGILDMMQSILIPNDLLEIYSSMQDI
CDRQKP010002909.1_Symbiodinium_sp.                      ...........................................................
JAUKPS010110286.1_Chlorellidium_tetrabotrys             ...........................................................
XP_004364654.1_Capsaspora_owczarzaki                    .....................................RMKQVKSLDPRVQALLDETVDE
KJE91799.1_Capsaspora_owczarzaki                        ..................................PTLMAKLAGPLHEAVQCELDALAGE
XP_048575736.1_Nematostella_vectensis                   .....................................DKEASKGPNLEILRCIYFIIRF
A0A6P8J3J2_Actinia_tenebrosa                            .....................................SKEATKQPNLEILRCIYFLIRF
XP_022780642.1_Stylophora_pistillata                    .....................................DNNASKTPNPELLRCIFFLFDF
A0A9X0CFV3_Desmophyllum_pertusum                        .....................................DKDASKQPNPELLRCIFFLIHF
A0A6S7HLM0_Paramuricea_clavata                          ........................................YAGQTNPELLRCFCFVLEF
XP_046856289.1_Xenia_sp.                                .....................................SMLYTTQCNLELLRCFRFIIEF
A0A813MIX4_Adineta_steineri                             ...........................................................
UJR19038.1_Adineta_vaga                                 GCVPSENILLDEYWKNAFSLHTSDENGL.FKRYFDSLNPQDSQNWPEDFNTLCRKFESF
JAXIUW010014154.1_Cephalothrix_simula                   ...............................................EIKCLERFLMDV
A0A8B8DK01_Crassostrea_virginica                        ..VP...........................................EIQRFLQFMKEK
XP_070564142.1_Ptychodera_flava                         ...............................................EIKRFFSFLSNF
ACQM01009845.1_Saccoglossus_kowalevskii                 .............................................NIEVNRFIMFLSNE
XP_078582628.1_Branchiostoma_floridae_japonicum         .......................................HGQAEIDQEIKDLVIKVLDF
JALCYT010000124.1_Branchiostoma_belcheri                .........................................QEEIDSEIKALADKVLSF
KAJ3014353.1_Thoreauomyces_humboldtii                   .....................................IVAEMTNVPNEVQEAVNSLFGW
KAJ3188706.1_Gaertneriomyces_sp._JEL0708                .....................................IVRYMRTVPDEVVALALDLFDW
XP_002110806.1_Trichoplax_adhaerens                     .........LVMFYKETFKNI...ENCKNMIRF....LQSQNFLIPSDLVEVLKELQTS
RDD37009.1_Trichoplax_sp._H2                            .........LVTFYTAIFPSLP.SQNSK.MINF....LKRSRFLIPLDLIDIFKSFQRP
A0A8W8L770_Magallana_gigas                              ..VS...........................................EIQRFIRFMEKE
```

*A0A8W8L770_Magallana_gigas*

*E8PLM2_Thermus_scotoductus*

```
E8PLM2_Thermus_scotoductus  ...........................................................
UJG41740.1_Candidatus_Heimdallarchaeum_aukensis  ...........................................................
MFW9996705_Candidatus_Odinarchaeota_archaeon  RYLTKLPAIK.................................................
MCH8915407.1_Thaumarchaeota_archaeon  ...........................................................
MHA1972560.1_Candidatus_Hodarchaeales_archaeon  ...........................................................
NHI95076.1_Candidatus_Lokiarchaeota_archaeon  ...........................................................
MEM2144305.1_Candidatus_Jordarchaeaceae_archaeon  CKKPHQI....................................................
AHB41284.1_candidate_division_SR1_bacterium  ...........................................................
HEV2339438.1_Patescibacteria_group_bacterium  ...........................................................
MBL9158386.1_Verrucomicrobiales_bacterium  ATPTQAVRPRRRRMAKVAAPKKGVTG.......ATKKAPLKAPRK..............
MGB8583703.1_Candidatus_Sulfotelmatobacter_sp.  VRQTFRGSPQE................................................
MBI3627515.1_Candidatus_Sungbacteria_bacterium  ...........................................................
_213105201.1_Candidatus_Protochlamydia_amoebophila  VEEKEIESKAEENSGTISFFAKVKNDIA..KNKQRKEEELKHKQAA.............
CDRQKP010002909.1_Symbiodinium_sp.  ...........................................................
JAUKPS010110286.1_Chlorellidium_tetrabotrys  ...........................................................
XP_004364654.1_Capsaspora_owczarzaki  VLKCAMA....................................................
KJE91799.1_Capsaspora_owczarzaki  MIARLVACRAS................................................
XP_048575736.1_Nematostella_vectensis  FKETLEKI...................................................
A0A6P8J3J2_Actinia_tenebrosa  FRETLQFK...................................................
XP_022780642.1_Stylophora_pistillata  FSQSTMNS...................................................
A0A9X0CFV3_Desmophyllum_pertusum  FTERLTAN...................................................
A0A6S7HLM0_Paramuricea_clavata  LQAV.......................................................
XP_046856289.1_Xenia_sp.  LNEK.......................................................
A0A813MIX4_Adineta_steineri  ...........................................................
UJR19038.1_Adineta_vaga  VKT........................................................
JAXIUW010014154.1_Cephalothrix_simula  DDKMDIDS...................................................
A0A8B8DK01_Crassostrea_virginica  H..........................................................
XP_070564142.1_Ptychodera_flava  KH.........................................................
ACQM01009845.1_Saccoglossus_kowalevskii  ASRI.......................................................
XP_078582628.1_Branchiostoma_floridae_japonicum  FNIPTDTATPPSGD..TASASSAQE..........DTSALSEQGTE.............
JALCYT010000124.1_Branchiostoma_belcheri  FNIQTAAGSPLSGQGTETPAGSGNETPA.....A.EGSALSGQGTE.............
KAJ3014353.1_Thoreauomyces_humboldtii  AQV........................................................
KAJ3188706.1_Gaertneriomyces_sp._JEL0708  AGV........................................................
XP_002110806.1_Trichoplax_adhaerens  YISV.......................................................
RDD37009.1_Trichoplax_sp._H2  DIELQRQNVKGQEKDKIEQRKGSQSQQRQFKELQDKNDSLRKSNESLEKKMEILTQCLN
A0A8W8L770_Magallana_gigas  ENKR.......................................................
A0A8W8L770_Magallana_gigas      .......................................................
```

*E8PLM2_Thermus_scotoductus*

```
E8PLM2_Thermus_scotoductus  .................................
UJG41740.1_Candidatus_Heimdallarchaeum_aukensis  ................................N
MFW9996705_Candidatus_Odinarchaeota_archaeon  .................................
MCH8915407.1_Thaumarchaeota_archaeon  .................................
MHA1972560.1_Candidatus_Hodarchaeales_archaeon  ................................N
NHI95076.1_Candidatus_Lokiarchaeota_archaeon  .................................
MEM2144305.1_Candidatus_Jordarchaeaceae_archaeon  .................................
AHB41284.1_candidate_division_SR1_bacterium  .................................
HEV2339438.1_Patescibacteria_group_bacterium  .................................
MBL9158386.1_Verrucomicrobiales_bacterium  ...............................RF
MGB8583703.1_Candidatus_Sulfotelmatobacter_sp.  .................................
MBI3627515.1_Candidatus_Sungbacteria_bacterium  .................................
_213105201.1_Candidatus_Protochlamydia_amoebophila  ......EDREKEDINEDFLVSSFFGSVDDSEDT
CDRQKP010002909.1_Symbiodinium_sp.  .................................
JAUKPS010110286.1_Chlorellidium_tetrabotrys  .................................
XP_004364654.1_Capsaspora_owczarzaki  .................................
KJE91799.1_Capsaspora_owczarzaki  ................................A
XP_048575736.1_Nematostella_vectensis  ................................S
A0A6P8J3J2_Actinia_tenebrosa  ................................I
XP_022780642.1_Stylophora_pistillata  .................................
A0A9X0CFV3_Desmophyllum_pertusum  .................................
A0A6S7HLM0_Paramuricea_clavata  ................................R
XP_046856289.1_Xenia_sp.  ................................H
A0A813MIX4_Adineta_steineri  .................................
UJR19038.1_Adineta_vaga  .................................
JAXIUW010014154.1_Cephalothrix_simula  ................................D
A0A8B8DK01_Crassostrea_virginica  .................................
XP_070564142.1_Ptychodera_flava  .................................
ACQM01009845.1_Saccoglossus_kowalevskii  .................................
XP_078582628.1_Branchiostoma_floridae_japonicum  ......NPY..PDL..................D
JALCYT010000124.1_Branchiostoma_belcheri  ......RAPEDPDL..................D
KAJ3014353.1_Thoreauomyces_humboldtii  .................................
KAJ3188706.1_Gaertneriomyces_sp._JEL0708  .................................
XP_002110806.1_Trichoplax_adhaerens  .................................
RDD37009.1_Trichoplax_sp._H2  KIEQEMESEECIDLRENLL..RHISEAREKVEK
A0A8W8L770_Magallana_gigas  ..............................FNN
A0A8W8L770_Magallana_gigas  ..............................
```
