## Supplemental tables_Figs for "Certain Aquatic Eukaryotes Harbor an OLD-Like Immune Defense System": Figure S2.pdf

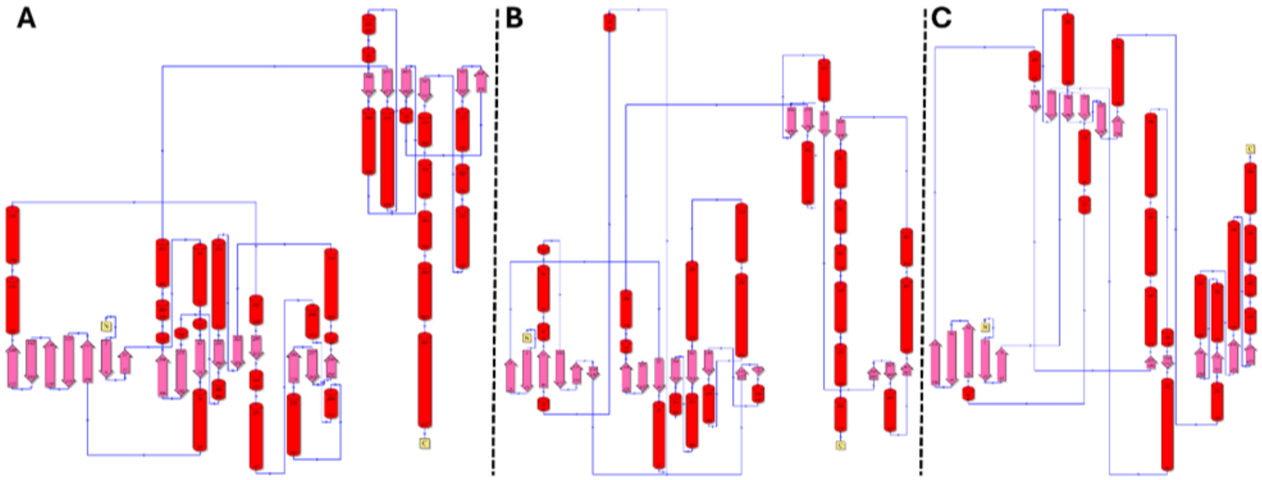


**Fig. S2. Topology diagrams**. (A) *M. gigas*, (B) *B. cereus* and (C) *T. scotoductus*. β-strands and α-helices are represented as pink arrows and red cylinders, respectively. N-terminal and C-terminal of sequences are represented by N and C letters in yellow, respectively.
