## Supplemental tables_Figs for "Certain Aquatic Eukaryotes Harbor an OLD-Like Immune Defense System": Figure S3.pdf

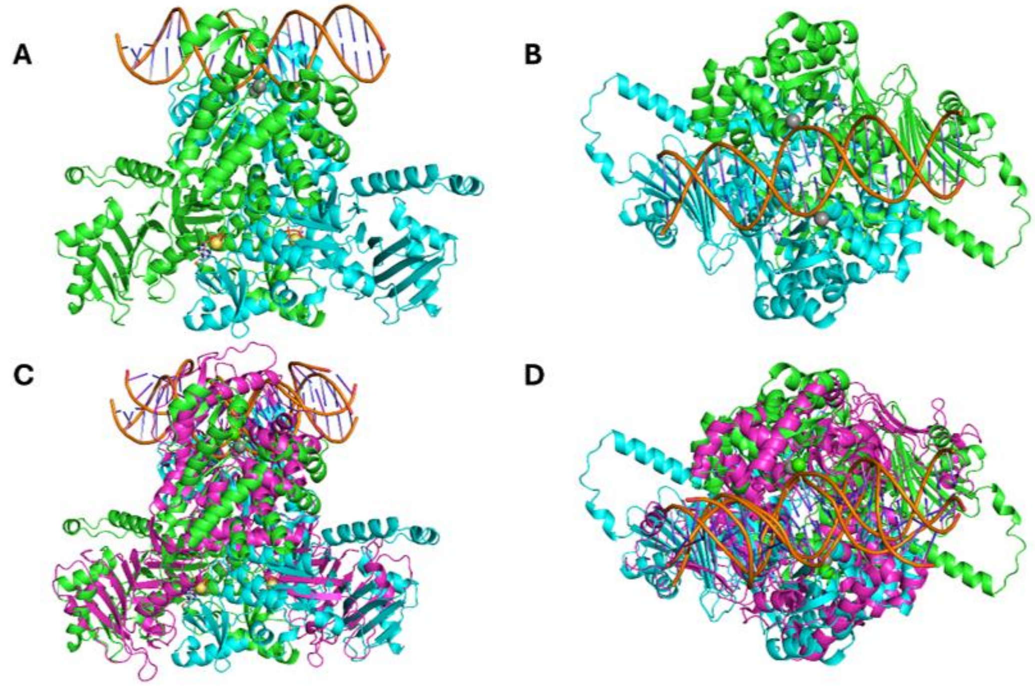


**Fig. S3. Predicted homodimer models of the *M. gigas* OLD-like protein bound to $Ca^{2+}$, $Mg^{2+}$, ATP, and dsDNA.** (**A**) Front and (**B**) top views of the *M. gigas* homodimer complex. Monomer A is shown in green, monomer B in cyan, dsDNA in orange, and ATP in light gray. $Ca^{2+}$ and $Mg^{2+}$ ions are represented as gray and yellow-orange spheres, respectively. (**C**, **D**) Structural superposition of the *M. gigas* homodimer-dsDNA with the *B. cereus* GajA homodimer (in pink) in complex with dsDNA and $Ca^{2+}$ (PDB ID: 8X51). Note the close spatial alignment and overlapping paths of the bound dsDNA molecules.
